# Treatment of a genetic liver disease in mice through transient prime editor expression

**DOI:** 10.1101/2024.01.22.575834

**Authors:** Tanja Rothgangl, Eleonora I. Ioannidi, Yanik Weber, András Tálas, Desirée Böck, Mai Matsushita, Elina Andrea Villiger, Lukas Schmidheini, Jennifer Moon, Paulo J.C. Lin, Steven H.Y. Fan, Kim F. Marquart, Cornelia Schwerdel, Nicole Rimann, Erica Faccin, Lukas Villiger, Hiromi Muramatsu, Máté Vadovics, Alessio Cremonesi, Beat Thöny, Manfred Kopf, Johannes Häberle, Norbert Pardi, Ying K. Tam, Gerald Schwank

## Abstract

Prime editing is a versatile genome editing technology that does not rely on DNA double-strand break formation and homology-directed repair (HDR). This makes it a promising tool for correcting pathogenic mutations in tissues consisting predominantly of postmitotic cells, such as the liver. While recent studies have already demonstrated proof-of-concept for *in vivo* prime editing, the use of viral delivery vectors resulted in prolonged prime editor (PE) expression, posing challenges for clinical application. Here, we developed an *in vivo* prime editing approach where we delivered the pegRNA using self-complementary adeno-associated viral (scAAV) vectors and the prime editor using nucleoside-modified mRNA encapsulated in lipid nanoparticles (LNPs). This methodology led to transient expression of the PE for 48h and 26% editing at the *Dnmt1* locus using AAV doses of 2.5×10^13^ vector genomes (vg)/kg and a single dose of 3mg/kg mRNA-LNP. When targeting the pathogenic mutation in the Pah*^enu2^* mouse model of phenylketonuria (PKU), we achieved 4.3% gene correction using an AAV dose of 2.5×10^13^ vg/kg and three doses of 2 mg/kg mRNA-LNP. Editing was specific to the liver and the intended locus, and was sufficient to reduce blood L-phenylalanine (Phe) levels from over 1500 µmol/l to below the therapeutic threshold of 600 µmol/l. Our study demonstrates the feasibility of *in vivo* gene correction in the liver with transient PE expression, bringing prime editing closer to clinical application.

## Introduction

Phenylketonuria (PKU) is an autosomal recessive metabolic liver disease caused by mutations in the phenylalanine hydroxylase (*PAH*) gene. While untreated PKU causes severe retardation, microcephaly and seizures, newborn screening followed by dietary L-phenylalanine (Phe) restriction and enzyme therapy leads to a life expectancy comparable to healthy individuals^1–4^. Nevertheless, despite existing treatments, learning disabilities and attention deficits remain frequent in PKU patients. In addition, the intricate dietary guidelines place a substantial burden on the quality of life. As a result, new treatment strategies attempting to permanently restore *PAH* expression in the liver are under exploration. Despite classical gene addition therapies, which provide an additional functional *PAH* gene copy to hepatocytes^5^, in the recent years there has been a growing interest in genome editing techniques that aim to directly repair pathogenic variants. The primary benefit of genome editing lies in the continuous expression of the corrected *PAH* allele, even as hepatocytes undergo cell division and genome replication. Hence, there is no need to express the genome editing tool over an extended period of time.

Employing the Pah*^enu2^* mouse model for human PKU, which contains a point mutation (c.835 T > C; p.F263S) that abolishes *PAH* function and results in Phe levels exceeding 1’500 µmol/l^6,7^, Richards et al. explored the feasibility of correcting this metabolic liver disease using CRISPR-Cas9 nucleases^8^. However, due to the low homology-directed repair (HDR) frequency in the liver only 1% of hepatocytes were corrected, which proved insufficient to resolve hyperphenylalaninemia. Circumventing the need for HDR, we and others have previously employed base editing to repair pathogenic PKU mutations at rates that led to therapeutic reduction of Phe levels^9–13^. Nonetheless, even though base editing holds promise for clinical use in PKU patients, the technology is largely limited to the correction of transition point mutations, excluding patients with different types of mutations.

Similar to base editing, prime editing allows precise correction of mutations without the need for HDR. Prime editors (PEs) consist of a H840A *Sp*Cas9 nickase (nCas9) fused to an engineered M-MLV reverse transcriptase (RT) (hereafter referred as PE2)^14^. This complex is guided to the locus of interest by the prime editor guide RNA (pegRNA), which contains a RT template (RTT) and a primer binding site (PBS) fused to the 3’ end of the guide RNA scaffold. nCas9-mediated nicking of the non-target DNA strand and hybridization of the PBS allows the RT to elongate the 3’ end using the RTT sequence as a template. Successful incorporation of the generated 3’ flap into the locus results in the installation of the intended edit. This mechanism therefore permits the introduction of any small-sized genetic change.

Previous studies established proof-of-concept for *in vivo* prime editing in the liver^15–17,18–20^. However, these studies employed viral delivery vectors, which resulted in permanent PE expression. This could pose challenges for clinical applications, as prolonged PE expression elevates the likelihood of installing unintended off-target mutations and potentially triggers T cell mediated elimination of edited cells that continue to express the genome editor. In this study, we established an *in vivo* prime editing approach where the pegRNA is delivered using self-complementary adeno associated virus (scAAV) vectors and the PE is delivered as mRNA encapsulated in lipid nanoparticles (LNPs). Transient prime editor expression resulted in editing rates of 26% at the *Dnmt1* locus and 4.3% at the *Pah^enu2^* locus, leading to therapeutically relevant reduction of Phe levels.

## Results

### Correction of the Pah^enu2^ mutation using optimized PE components and AAV delivery

In a previous study we attempted to correct the pathogenic mutation in the *Pah^enu2^* mouse model for PKU using AAV-mediated delivery of intein-split PE2 and a pegRNA targeting *Pah^enu2^* (mPKU-2.1)^19^. However, despite applying AAV doses of 1x 10^14^ vector genomes (vg)/kg correction rates remained below 1%, and treatment was only successful when prime editing components were delivered from adenoviral (AdV) vectors at doses that are not viable in a clinical context. This prompted us to test if recent improvements in the prime editing technology could lead to an increase in correction rates of the *Pah^enu2^* mutation.

We exchanged PE2 with PEmax^21^, a prime editor variant with optimized codon usage and NLS-linker design, and incorporated pseudoknot structures to the 3’ end of the pegRNA to protect them from exonuclease degradation (epegRNAs)^22,23^. Additionally, we modified the RTT sequence of the pegRNA to co-introduce a silent mutation in the GG sequence of the PAM, preventing retargeting of the locus once the edit is installed (**Fig. S1b**). Importantly, when transfected into HEK293T cells with the murine *Pah^enu2^* locus stably integrated (**Fig. S1a**), the adapted editing components led to substantially higher correction rates without inducing indel mutations above background (**Fig. 1a-c**).

**Figure 1.**
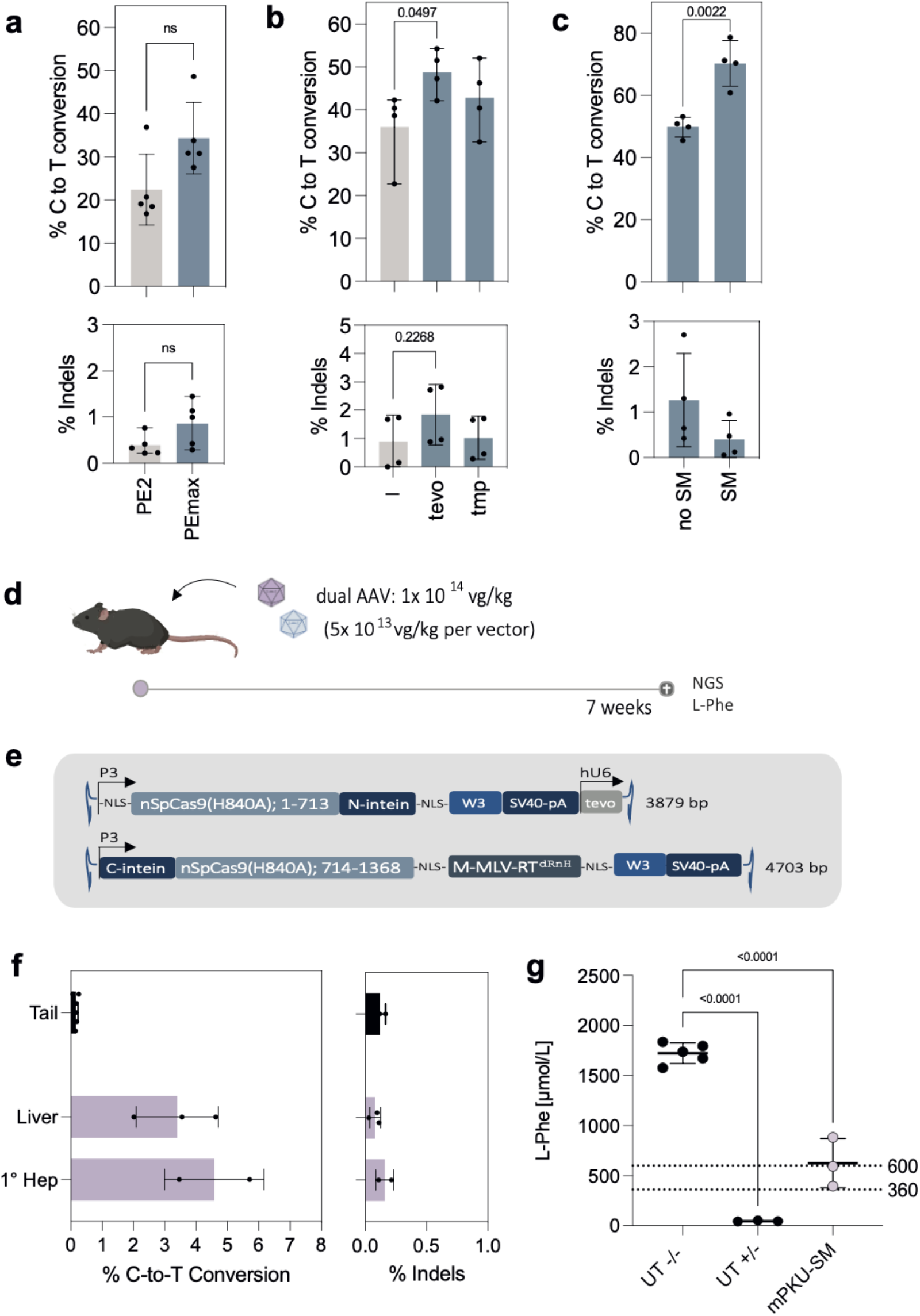
*In vivo* correction of *Pah^enu2^* mice using AAV-mediated delivery of intein-split PE. (**a**) C-to-T conversion rates (upper panel) and indel rates (lower panel) in HEK293T cells with the integrated *Pah^enu2^* locus, transfected with PE2 vs. PEmax, or with (**b**) unmodified (gray) vs. tevopreQ_1_ (tevo) modified vs. tmpknot (tmp) modified pegRNAs (Suppl. Table 1), or with (**c**) tevopreQ1-mPKU-2.1 vs. tevopreQ1-mPKU-SM. Black arrowheads indicate the site of the silent mismatch (SM) in the reverse transcriptase template (RTT). The part of the RTT complementary to the PAM is highlighted in grey. Values represent mean +/- s.d. of four independent biological replicates. Means were compared using an unpaired Student’s t tests. (ns, not significant, P > 0.05). (**d**) Schematic illustration of the experimental outline. Adult *Pah^enu2^* mice were injected with the dual AAV system. After 7 weeks blood was taken to analyze Phe levels, and whole liver lysates and isolated hepatocytes were analyzed by NGS to assess editing rates. (**e)** Schematic illustration of the AAV constructs selected for *in vivo* experiments. The tevopreQ_1_-mPKU-SM pegRNA was expressed from a hU6 promoter on the N-terminal vector. Indicated are AAV genome lengths including ITRs in base pairs (bp). (**f**) C-to-T conversion rates *in vivo* in PKU mice. Animals (n=3) were treated with an AAV dose of 1×10^14^ vg/kg (5×10^13^ vg/kg per vector). Editing rates were assessed in lysed tail tissue (n=3), liver tissue (n=3) and isolated hepatocytes (n=3 and n=2). (**g**) Phe levels at the experimental endpoint of untreated homozygous *Pah^enu2^* animals (UT -/-), untreated heterozygous animals (UT +/-) and homozygous animals treated with prime editing. Dashed lines indicate recommended therapeutic thresholds for Phe levels in adults (600 μmol/L) or in children/during pregnancy (360 μmol/L)^28,29^. Values represent mean +/- s.e.m of independent biological replicates and were analyzed using an ordinary one-way ANOVA using Dunnett’s multiple comparisons test. W3, truncated version of the posttranscriptional regulatory element of woodchuck hepatitis virus (WPRE); nSpCas9, nickase of Streptococcus pyogenes Cas9; tevo, trimmed engineered pegRNA; M-MLV-RT^dRnH^, M-MLV (Moloney Murine Leukemia Virus) Reverse Transcriptase (RT)-delta RnaseH; SV40-pA, Simian virus 40 poly-adenylation signal; NLS, nuclear localization signal. Unless otherwise stated, values depict mean +/- s.d. of independent biological replicates (ns, not significant, P > 0.05).

Next, we assembled AAV vectors for *in vivo* delivery of the optimized PE components (PEmax + tevopreQ_1_-mPKU-SM) into *Pah^enu2^* mice. Since the size of the PE exceeds the packaging capacity of AAV, we and others have previously employed the intein-split system to express PE2 from two separate AAVs^15,16,24^. Here, we tested two intein-split designs for PEmax in HEK293T cells (1153/1154 and 713/714). While both variants showed comparable activity (**Fig. S1c**), we selected the 713/714 variant for our *in vivo* experiments as it facilitates the generation of AAV constructs that are more equal in size when the non-essential RnaseH domain of the RT is removed and the pegRNA is positioned on the vector containing the N-intein. The tevopreQ_1_-mPKU-SM pegRNA was cloned downstream of the human U6 promoter, and N- and C-terminal fragments of PEmax were cloned between the liver-specific P3 promoter^25^ and the 3’ UTR, which contains the W3 post-transcriptional regulatory element^26^ and the simian virus (SV40)-polyA tail (**Fig. 1d, S1d)**. Recombinant AAV2 genomes were then packaged into hepatotropic AAV serotype 9 capsids (AAV2/9) and systemically administered to *Pah^enu2^* mice via the tail vein in a 1:1 ratio at a final dose of 1×10^14^ vg/kg (**Fig. 1e**). Analysis of isolated hepatocytes by next generation sequencing (NGS) after a period of 7 weeks revealed 4.6% correction of the *Pah^enu2^* mutation (**Fig. 1f**), which resulted in a reduction of blood Phe levels from over 1500 µmol/L to 623.7 µmol/L (**Fig. 1g; S2a**). In line with previous genome editing studies that utilized AAV9 vectors in combination with the hepatocyte-specific P3 promoter^7,25,27^, we found that editing was largely limited to hepatocytes (**Fig. S2b**). Furthermore, we did not detect unintended editing in the liver of treated mice at the top 5 off-target binding sites of the mPKU-SM pegRNA, which were previously identified by CHANGE-seq^19^ (**Fig. S2c,d**).

### In vivo prime editing using LNP mediated pegRNA and PE mRNA delivery

While *Pah^enu2^* correction rates with optimized PE components were substantially increased compared to our initial study^22^, they were not sufficient to reduce Phe levels below the therapeutic threshold for adult PKU patients (600 µmol/L)^28^. In addition, recent clinical trials revealed that AAV doses of 1×10^14^ vg/kg could lead to severe immune reactions and, in rare incidents, have been associated with patient mortality^30,31^. Finally, AAV-mediated delivery leads to prolonged PE expression, which is not desired for genome editing as it could lead to an accumulation of off-target mutations and induce an immune response to the bacterial Cas9 or the viral RT.

Previously, lipid nanoparticle (LNP)-mediated mRNA and sgRNA delivery has been employed for transient genome editing with Cas9 nucleases and base editors^7,27,32–36^. To assess if a similar approach is feasible for prime editing, we generated nucleoside-modified mRNAs^37,38^ encoding for PE2 and PEmax and packaged them into LNPs. Confirming transient liver expression after systemic delivery of 2 mg/kg LNP-mRNA, we observed a peak in PE mRNA levels at 6 hours post injection (h.p.i.) and a peak in protein levels at 24 h.p.i., while at 45 h.p.i, neither PE mRNA nor PE protein levels were detectable anymore (**Fig. 2a,b; Fig. S3a,b**). Next, we chemically synthesized the tevopreQ_1_-mPKU-SM pegRNA targeting the *Pah^enu2^*locus, and a pegRNA that installs a G-to-C edit at the *Dnmt1* locus. This pegRNA has previously been successfully utilized for *in vivo* prime editing in the liver using AAV and AdV mediated delivery^19^. To protect pegRNAs from exonucleases they were modified with 2′-O-methyl-3′-phosphorothioate (MS) at the 5’ end and 2′-O-methyl-3′-phosphonoacetate (MP) at the 3’ end^39^. In addition, we generated pegRNAs in which the RTT was substituted with deoxyribose nucleotides to further protect it from degradation by endonucleases (DNA-mod pegRNAs) (**Fig. 2c**). Confirming functionality of the synthesized pegRNAs, co-electroporation with PEmax mRNA into HEK reporter cells resulted in 25% editing at the *Pah^enu2^*site and 37% editing at the *Dnmt1* site using non-DNA modified pegRNAs, and 3% editing at the *Pah^enu2^* site and 21% editing at the *Dnmt1* site using DNA-modified pegRNAs (**Fig. 2d**). However, when administered into mice, using a repeated dosing scheme of 2 mg/kg LNP containing PEmax mRNA followed by 2 mg/kg LNP containing the respective pegRNA (to assure presence of PE protein when the pegRNA is delivered; **Fig. 2e**), editing rates remained below 2% at the *Dnmt1* locus and 1% at the *Pah^enu2^* locus (**Fig. 2f**).

**Figure 2.**
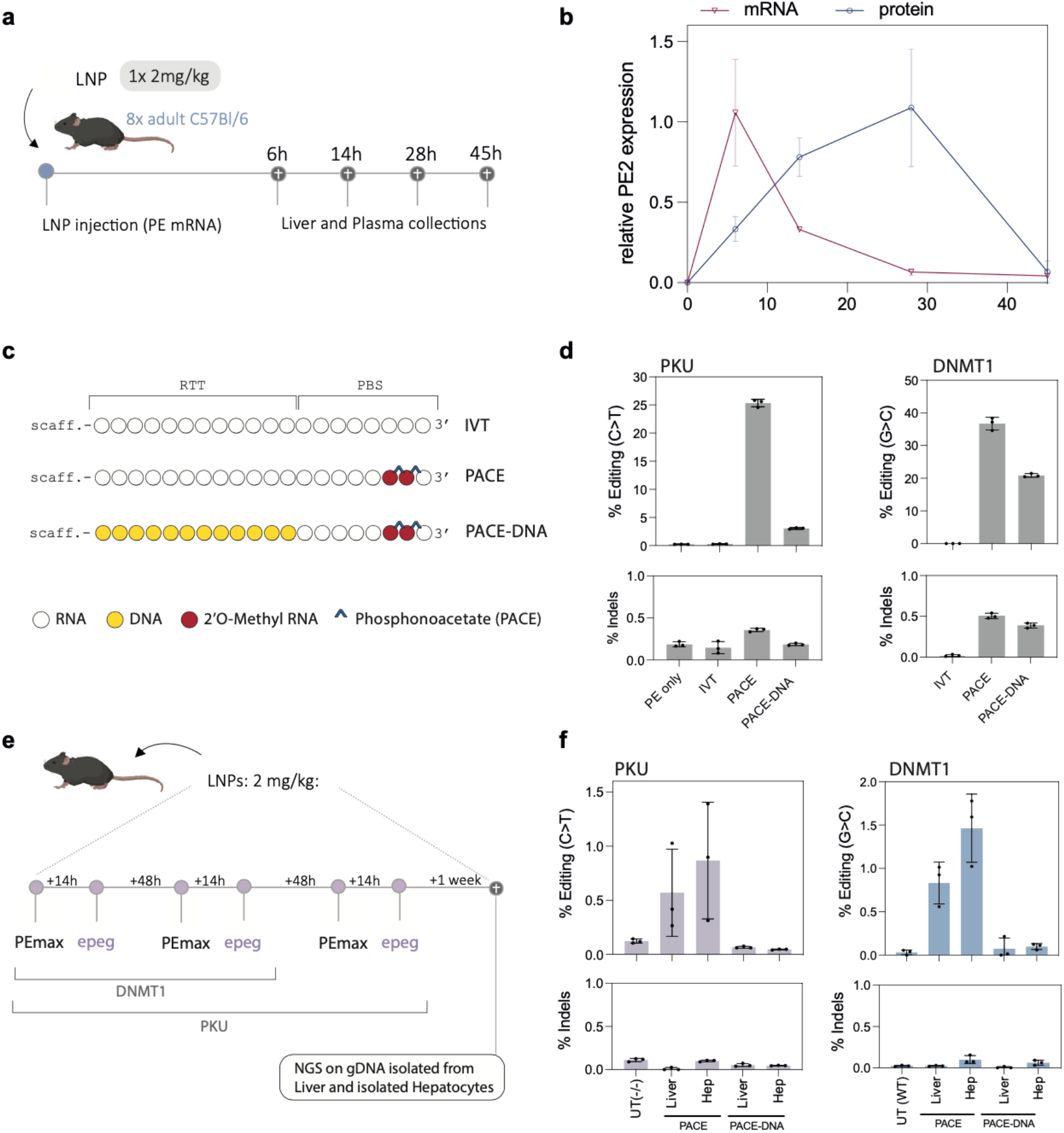
In vivo correction of Pah*enu2* mice using LNP-RNA delivery. (**a**) Schematic illustration of the experimental setup. C57Bl/6J animals were injected with LNP-mRNA (2 mg/kg) encoding for PEmax. Two animals were sacrificed at each experimental endpoint (6, 14, 28 and 45 hours post injection; h.p.i.) and liver tissue was isolated for downstream processing. (**b**) Expression kinetics of prime editor mRNA and protein levels. Relative values were normalized to the respective housekeeping gene and the average observed peak expression (6 and 28 h.p.i. respectively). Each value represents the mean +/- s.e.m. of two individual biological replicates. (**c**) Illustration of the employed pegRNAs. Full pegRNAs sequences are listed in Suppl. Table 4. (**d**) *In vitro* editing rates of the *Pah^enu2^* locus (left panel) and *Dnmt1* locus (right panel) after electroporation of HEK293T reporter cells with the differently modified pegRNAs depicted in (c). **(e)** Schematic illustration of the experimental setup. LNP containing PEmax mRNA was injected 14 hours before LNP-pegRNA was delivered into *Pah^enu2^* or C57BL/6J mice at a dose of 2 mg/kg. *Pah^enu2^* mice were redosed three times while C57BL/6J mice were redosed twice. At the experimental endpoint, NGS was performed on isolated hepatocytes and whole liver lysates. (**f)** *In vivo* editing rates of homozygous (-/-) *Pah^enu2^* animals (left panel) and wildtype (WT) C57BL/6J animals (right panel). Unless otherwise stated, values represent mean +/- s.d. of independent biological replicates.

### In vivo correction of adult Pah^enu2^ mice using AAV-pegRNA and LNP-mRNA delivery

We reasoned that the primary constraint for prime editing efficiency could be the abundance of functional pegRNA rather than the PE. Therefore, we also tested an alternative strategy, where only the PE mRNA is delivered via LNP and the pegRNA is delivered via AAV. PKU mice were first administered with 2.5×10^13^ vg/kg self-complementary AAV2/9 (scAAV2/9) encoding for the tevopreQ_1_-mPKU-SM pegRNA (**Fig S4a**), followed by treatment with PE2 LNP-mRNA or PEmax LNP-mRNA at a dose of 3 mg/kg (**Fig 3a**). In addition, we injected wildtype mice with LNP-mRNA that were pre-treated with an scAAV2/9 encoding for the tevopreQ1-modified *Dnmt1* pegRNA (**Fig 3a**). While editing rates were relatively low at *Pah^enu2^* locus (1.1% with PE2 and 1.0% with PEmax) (**Fig. 3b**), at the *Dnmt1* locus we observed 17.7% editing with PE2 and 26.2% editing with PEmax (**Fig. 3c**). Notably, editing rates were similar when the scAAV2/9 dose was decreased from 2.5 x 10^13^ vg/kg to 1 x 10^12^ vg/kg (20% vs. 26%; **Fig. 3d**), and when animals were analyzed 4 months after treatment instead of 1 week after treatment (**Fig. S4b**).

**Figure 3.**
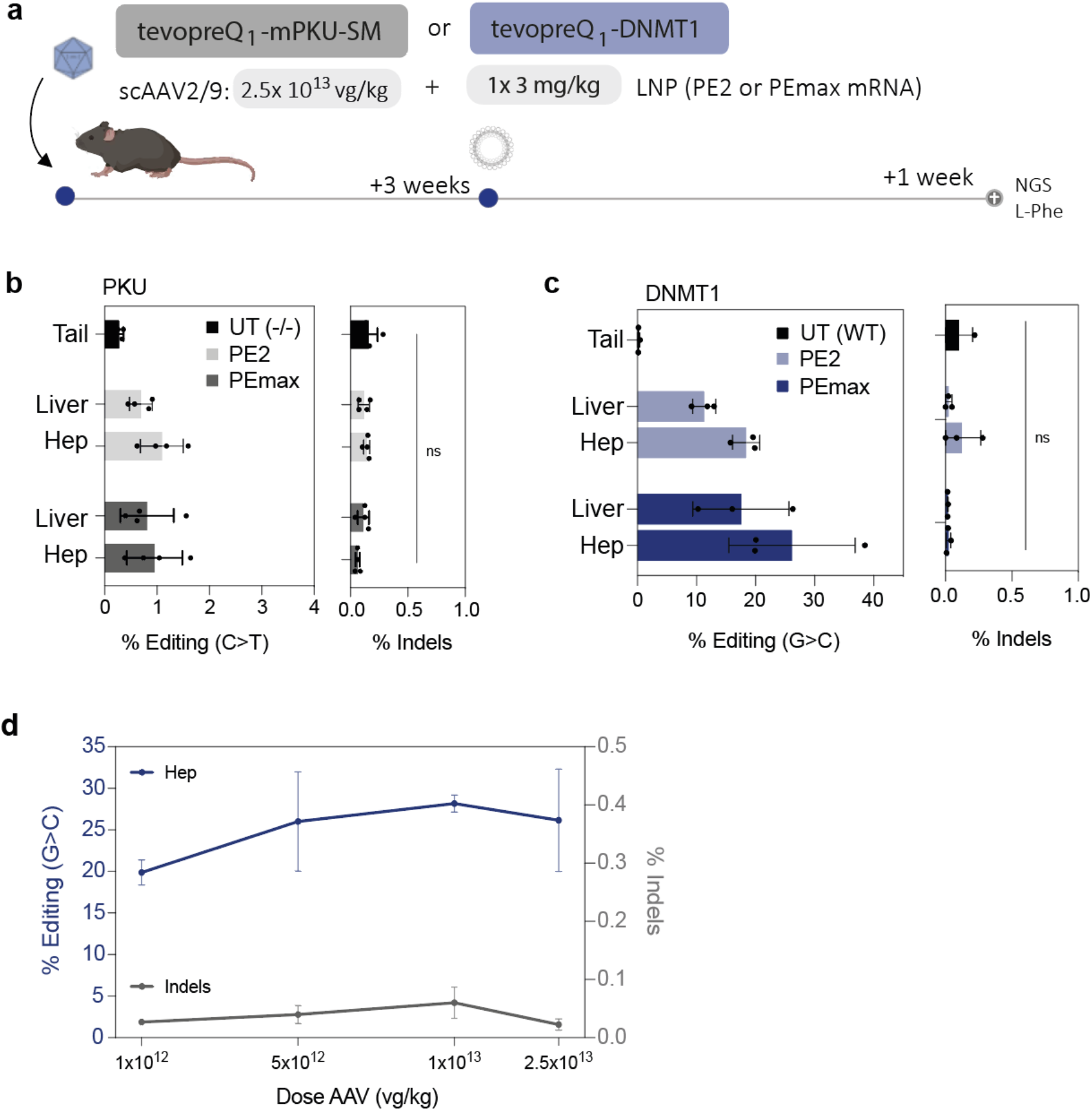
*In vivo* prime editing using AAV-pegRNA and LNP-mRNA delivery. (**a**) Schematic illustration of the experimental setup. scAAV2/9 expressing the tevopreQ_1_-Dnmt1 pegRNA or tevopreQ1-mPKU-SM was delivered into C57BL/6J mice or *Pah^enu2^* mice, respectively, at a dose of 2.5×10^13^ vg/kg. Mice were subsequently injected with LNP (one time 3 mg/kg) containing PE2 or PEmax mRNA. At the experimental endpoint, NGS was performed in isolated hepatocytes, whole liver lysates. (**b**) *Pah^enu2^* correction rates (left panel) and indel rates (right panel) of untreated tail tissue (n=5) or animals treated with a single dose of 3mg/kg PE2 (n=4) or PEmax (n=4) mRNA-LNP. (**c**) G-to-C editing rates (left panel) and indel rates (right panel) at the *Dnmt1* locus in untreated tail tissue (n=3) or animals treated with a single dose of 3mg/kg PE2 (n=3) or PEmax (n=3) mRNA-LNP. Values represent mean +/- s.d. of independent biological replicates and were analyzed using unpaired Student’s t tests (ns, not significant, P > 0.05) (**d**) G-to-C editing rates (blue line, left y-axis) and indel rates (grey bar, right y-axis) of animals treated with one dose of 3 mg/kg PEmax LNP-mRNA and pre-treatment of different doses of scAAV encoding for the *Dnmt1-*targeting pegRNA. Values represent mean +/- s.e.m. of independent biological replicates.

Since correction rates at the *Pah^enu2^* locus with the dual AAV-pegRNA and LNP-mRNA approach were not sufficient to substantially reduce Phe levels, we next dosed AAV-pegRNA treated animals three times with 2 mg/kg mRNA-LNP in a 48-hour interval (**Fig 4a**). Importantly, the redosing regimen increased editing rates up to 6.2% with PE2 (average 2.9%) and up to 8.7% with PEmax (average 4.3%; **Fig. 4b)**. This led to a reduction of blood L-Phe to levels below the therapeutic threshold of 600 µmol/L (**Fig. 4c,d**), and a corresponding increase in *Pah* enzyme activity in the liver (**Fig. 4e)**. Moreover, similar to the *Dnmt1* locus also at the *Pah^enu2^* locus editing rates did not significantly decrease when the scAAV2/9 dose was lowered from 2.5×10^13^ vg/kg to 5×10^12^ vg/kg (**Fig. S4c**).

**Figure 4.**
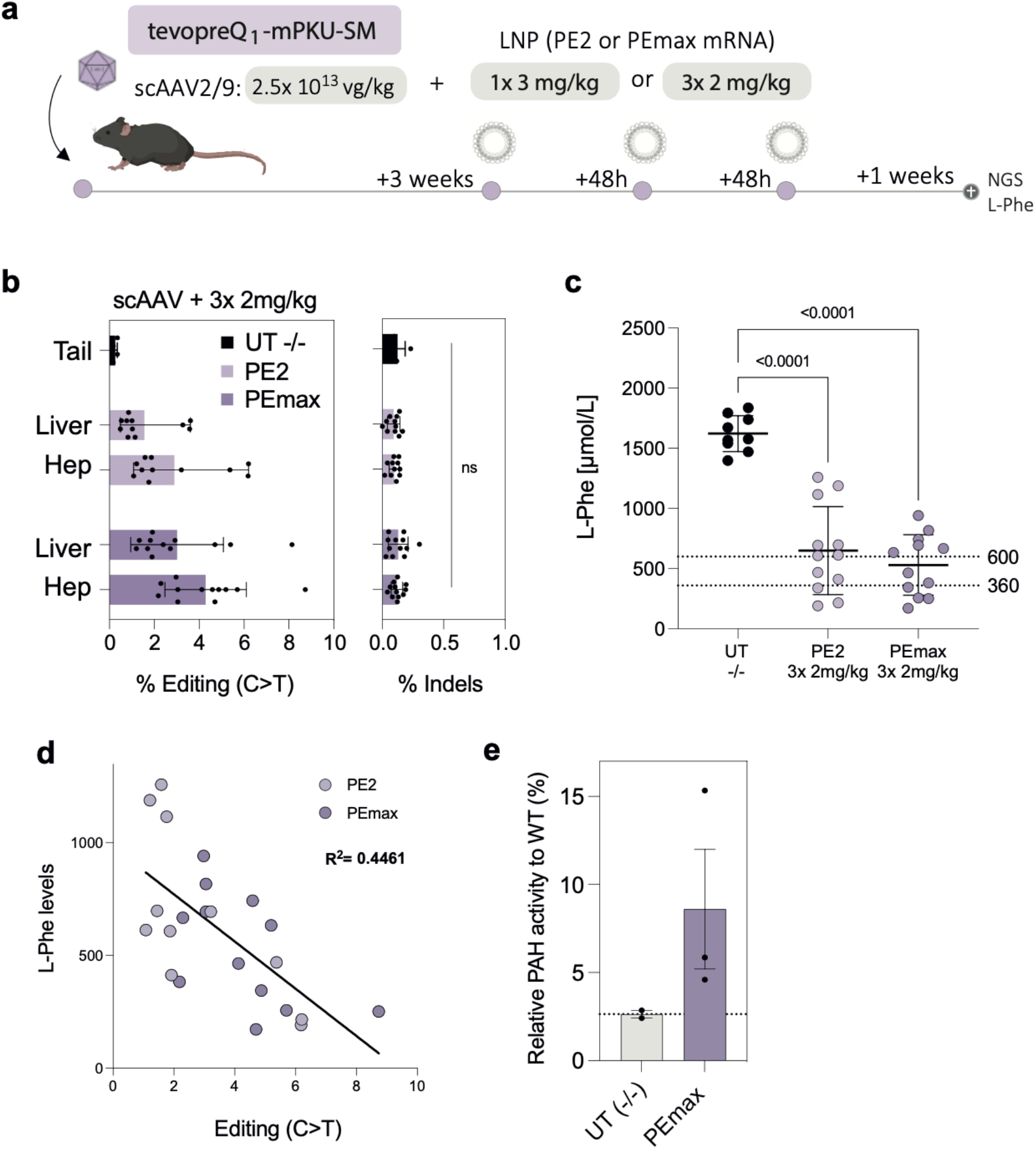
In vivo correction of *Pah^enu2^* mice using AAV-pegRNA and LNP-mRNA delivery. **(a)** Schematic illustration of the experimental setup. scAAV2/9 expressing the tevopreQ1-mPKU-SM pegRNA was delivered into *Pah^enu2^* mice at a dose of 2.5×1013 vg/kg, which were subsequently injected (one time 3 mg/kg or three times 2 mg/kg) with LNP containing PE2 or PEmax mRNA. At the experimental endpoint Phe levels were measured and NGS was performed in isolated hepatocytes, whole liver lysates and lysates from other organs. **(b)** *Pah^enu2^* correction rates (left panel) and indel rates (right panel) of untreated tail tissue (n=5) or animals treated with three doses of 2 mg/kg PE2 (n=12) or PEmax (n=12) mRNA-LNP. **(c)** Phe levels at the experimental endpoint of untreated homozygous *Pah^enu2^* animals (UT -/-; n=5), and homozygous Pahenu2 animals dosed three times with 2mg/kg PE2 (n=12) or PEmax (n=12). Dashed lines indicate recommended therapeutic thresholds for L-Phe levels in adults (600 μmol/L) or in children/during pregnancy (360 μmol/L)31,33. Values represent mean +/- s.e.m of independent biological replicates and were analyzed using an ordinary one-way ANOVA using a Dunnett’s multiple comparisons test. **(d)** Correlation between C-to-T editing rates and Phe levels in PEmax (dark magenta) and PE2 (light magenta) treated animals at the experimental endpoints. R^2^, coefficient of determination; Values represent mean +/- s.d. of independent biological replicates. **(e)** Enzyme activity of *Pah* in liver tissue lysates from untreated homozygous *Pah^enu2^* animals and homozygous *Pah^enu2^* animals treated with scAAV (pegRNA) and PEmax LNP-mRNA. Values are depicted as relative values to *Pah* activity in wt animals. Unless otherwise stated, values represent mean +/- s.d. of independent biological replicates and were analyzed using unpaired Student’s t tests (ns, not significant, P > 0.05)

### In vivo prime editing using AAV-pegRNA and LNP-mRNA did not result in off-target editing or liver damage

To assess if prime editing with AAV-pegRNA and LNP-PE mRNA delivery was limited to the liver, DNA of treated animals was isolated from other organs and analyzed by NGS. Consistent with previous mRNA-LNP biodistribution studies^40^, we did not observe substantial editing in any of the analyzed tissues in *Pah^enu2^*targeted animals treated with three doses of 2 mg/kg LNP-mRNA or *Dnmt1*-targeted animals treated with one dose of 3mg/kg LNP-mRNA (**Fig. 5a, S5a**). Next, we assessed if editing occurred at other sites in the genome and performed targeted amplicon sequencing at the top 5 off-target binding sites of both pegRNAs, which were previously identified by CHANGE-seq^19^ (**Fig. S2c, S5b**). Importantly, we did not observe editing above background in treated animals at any of the potential off-target sites (**Fig. 5b,c**). Finally, we examined whether delivery of LNP-mRNA or scAAV triggered liver toxicity or innate immune responses. While a slight elevation of alanine transaminase (ALT) and aspartate aminotransferase (AST) was observed between 6-28 h after administration of 2 mg/kg PEmax LNP-mRNA, levels returned to baseline levels at 45 h.p.i. (**Fig. 5d,e**). Likewise, the transient elevation of the proinflammatory cytokines Monocyte Chemoattractant Protein-1 (MCP-1), Interleukin-1 alpha (IL-1α), Interferon-gamma (IFNy) and Tumor necrosis factor alpha (TNF-α), which was observed at 6 h.p.i., was not detectable anymore at later timepoints (**Fig. 5f-i)**. Finally, also delivery of scAAV2/9 encoding for the pegRNA at a dose of 2.5 x 10^13^ vg/kg did not induce elevated levels of ALT, AST or any of the tested proinflammatory cytokines (**Fig. 5d-i**). In line with these observations, histological examination of the liver of treated animals did not reveal any obvious signs of tissue necrosis (**Fig. S5c**).

**Figure 5.**
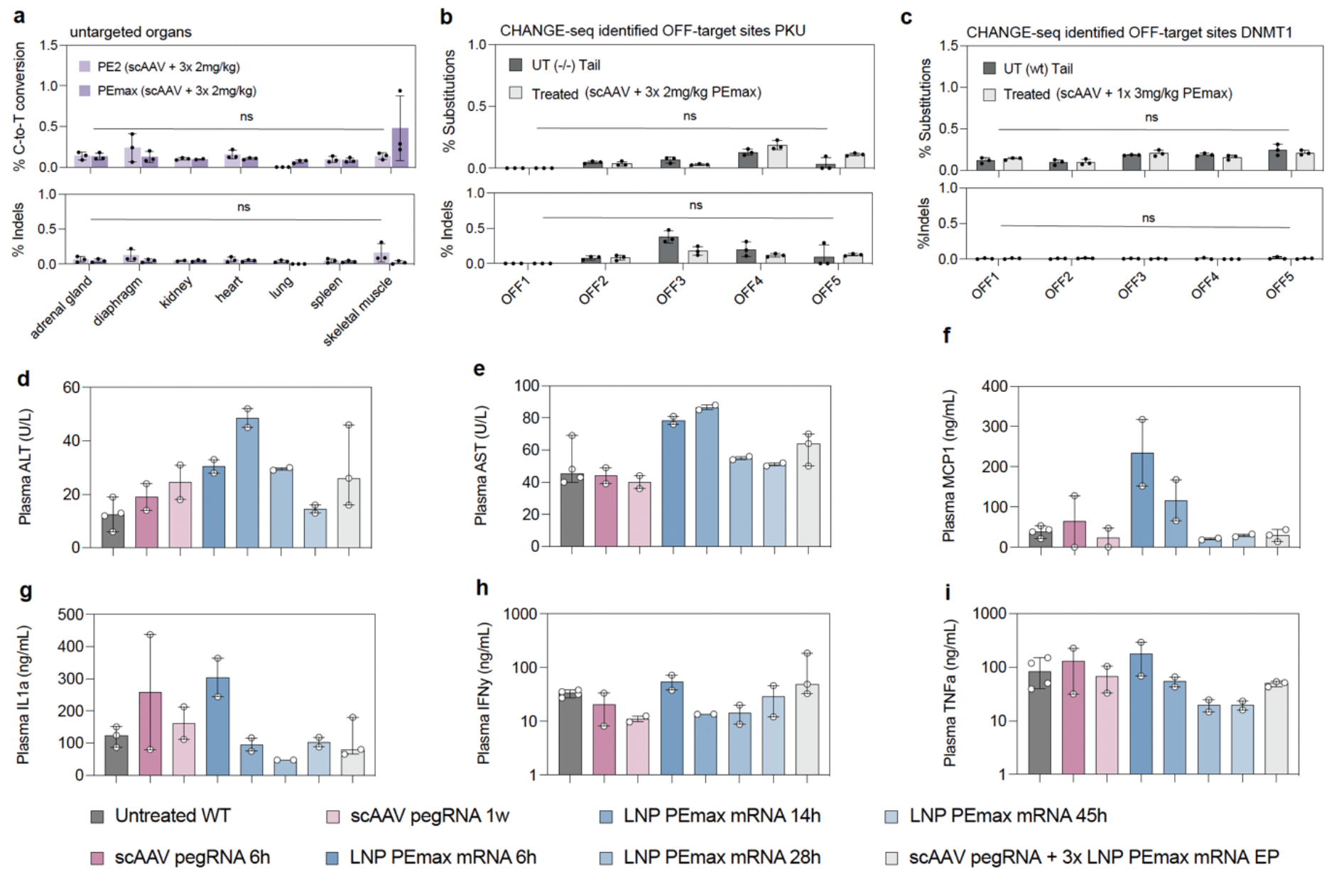
Assessment of off-target effects and liver toxicity in animals treated with AAV-pegRNA and LNP-mRNA. (**a**) C-to-T conversion rates (upper panel) and indel rates (lower panel) at the *Pah^enu2^* locus on DNA isolated from tissues other than the liver (n=3). Animals were pre-treated with scAAV encoding for the *Pah^enu2^* targeting pegRNA and dosed three times with 2 mg/kg PEmax. Editing rates were assessed by NGS on genomic DNA isolated from whole tissue lysates. (**b**) Indel rates and substitution rates at CHANGE-seq identified off-target sites for the *Pah^enu2^* targeting pegRNA^19^. Tail tissue (UT (-/-) Tail) was obtained from untreated animals and hepatocytes were isolated from animals treated with 3x 2mg/kg LNP-PEmax. (**c**) Indel rates and substitution rates of CHANGE-seq identified off-target site^19^ for the *Dnmt1*-targeting pegRNA. Tail tissue was obtained from untreated C57BL/6J mice (UT (wt) Tail) and hepatocytes were isolated from animals treated with 1x 3mg/kg LNP-PEmax. Values represent mean +/- s.d. of three independent biological replicates. **(d-i)** Measured concentration of the respective markers for liver damage or inflammation in the plasma of animals treated between 0-45 hours: Alanine aminotransferase (ALT) (d), aspartate aminotransferase (AST) (e), Monocyte Chemoattractant Protein-1 (MCP-1) (f), Interleukin-1 alpha (IL-1α) (g), Interferon-gamma (IFNy) (h) and Tumor necrosis factor alpha (TNF-α) (i); Values represent median +/- range of at least two independent biological replicates. Means were compared using Šídák’s multiple comparisons test (ns, not significant, P > 0.05).

## Discussion

Previous studies have demonstrated the feasibility of *in vivo* prime editing using AAV or AdV delivery vectors ^15,16,20,24,41–45^. While these studies highlight the potential of *in vivo* prime editing for correcting genetic diseases, prolonged PE expression from viral vectors may limit clinical application of these approaches. Similarly, previous studies achieved substantial reduction of Phe levels in PKU mice by delivering a functional copy of the *Pah* gene on an AAV vector^46^, but since AAV vectors remain episomal and hepatocytes divide approximately once every year^47^ it is unlikely that this approach could result in a lifelong cure of patients. Here, we developed an *in vivo* prime editing approach where only the pegRNA is delivered via AAV and the PE is delivered as LNP-mRNA. This approach led to transient PE expression in the liver, which resulted in 26% editing at the *Dnmt1* locus using a single dose of 3 mg/kg LNP-mRNA and 4.6% editing at the *Pah^enu2^* locus using three doses of 2mg/kg LNP-mRNA. Importantly, *Pah^enu2^* correction rates were sufficient to reduce Phe levels below 600 µmol/l, which is considered therapeutic for adult PKU patients. In line with the high specificity of prime editing described in previous studies^48^, we did not detect editing at experimentally validated off-target binding sites with both pegRNAs. Furthermore, hepatotropism of LNPs and AAV2/9 ensured that editing was largely restricted to hepatocytes.

The efficiency of prime editing is influenced by the sequence of the pegRNA and the target site, making it challenging to address all pathogenic PKU mutations effectively. However, transient PE expression using LNP-mediated mRNA delivery could prove to be a valuable strategy for treating certain patients with PKU or other genetic liver disorders.

## Materials and Methods

### Generation of plasmids

To generate epegRNA plasmids, annealed spacer, scaffold, and 3′ extension oligos were cloned into pU6-tevopreq1-GG-acceptor (Addgene No.174038) by Golden Gate Assembly as previously described^14^. For the generation of split-intein PEmax constructs, inserts were PCR amplified from the epegRNA plasmids and from pCMV-PEmax (Addgene No. 174820) and inserted into the respective backbones (Addgene No. 187181 and 117182) using HiFi DNA Assembly Master Mix [New England Biolabs (NEB)]. To generate PiggyBac PKU reporter plasmids, inserts with homology overhangs for cloning were ordered from IDT and cloned into the pPB-Zeocin backbone using HiFi DNA Assembly Master Mix (NEB). All PCRs were performed using Q5 High-Fidelity DNA Polymerase (NEB).

### Cell culture transfection and genomic DNA preparation

HEK293T [American Type Culture Collection (ATCC): CRL-3216] and K562 (ATCC: CCL-243) cells were maintained in Dulbecco’s modified Eagle’s medium (DMEM) plus GlutaMAX (Thermo Fisher Scientific) or Roswell Park Memorial Institute (RPMI) 1640 Medium (Thermo Fisher Scientific), respectively and supplemented with 10% (v/v) fetal bovine serum (FBS) and 1% penicillin/streptomycin (Thermo Fisher Scientific) at 37°C and 5% CO2. Cells were maintained at confluency below 90% (HEK293T) or below a density of 1.8 Mio. cells per milliliter (K562) and seeded into 48-well cell culture plates (Greiner). About 0.1 Mio. cells were transfected using 1.5 μl of Lipofectamine 2000 (Thermo Fisher Scientific) with 375 ng of PEmax and 125 ng of epegRNA or 250 ng of each AAV construct according to the manufacturer’s instructions. Unless otherwise noted, cells were incubated for 3 days, and genomic DNA was isolated by direct lysis. For nucleofections 0.5 pmol of the PEmax mRNA were used in a 1:1 molar ratio with the pegRNAs. Nucleofections of mRNA were carried out with one pulse of 1400 mV and 20 ms pulse width. After nucleofection, cells were cultured in 200 uL of DMEM plus GlutaMAX for 48 hours prior to isolation of genomic DNA by direct lysis. In vitro transcription of the unmodified pegRNA was carried out as previously described on a synthetic DNA fragment (Table S4)^49^.

### Generation of reporter cells by PiggyBac transposon

For generation of the PKU reporter cell lines with the PiggyBac transposon, 75’000 HEK293T or K562 cells were seeded into a 24-well culture plate (Greiner) and transfected the next day using Lipofectamine 2000 (Thermo Fisher Scientific) according to the manufacturer’s instructions. Briefly, 1.5 μl of Lipofectamine was mixed with 23.5 μl of Opti-MEM, incubated at room temperature for 10 min, and added to 225 ng of transposon plasmid and 25 ng of transposon helper plasmid (filled up to 25 μl of Opti-MEM). After 30 min of incubation at RT, cells were transfected. Three days after transfection, cells were enriched for 10 days using Zeocin (InvivoGen, 150 μg/ml).

### AAV production

Pseudotyped AAV9 vectors (AAV2/9) were produced by co-transfection of packaging (see above), capsid and helper plasmids (Addgene No. 112865 and 112867) were incubated for five days until harvest and precipitation using PEG and NaCl. The AAVs were further purified using a gradient centrifugation with OptiPrep (Sigma-Aldrich) as previously described^50^ and subsequently concentrated using Vivaspin® 20 centrifugal concentrators (VWR). Physical titers (vector genomes mL^−1^) were determined using a Qubit 3.0 Fluorometer. In brief, the Qubit Fluorometer 3.0 (Life Technologies) was used to measure the concentrations (ng/mL) of the extracted genomes by denaturation at 95°C for 5 min, after which the readings were converted to vector genomes per mL using the genome’s molecular mass and Avogadro’s constant. Identity of the packaged genomes of each AAV vector was confirmed by Sanger DNA sequencing by Mycrosynth AG (Balgach, Switzerland), testing 500 ng of denatured AAV using an AAV-genome-specific sequencing primer. AAV2/9 viruses were stored at −80°C until use and diluted with phosphate-buffered saline (PBS, Thermo Fisher Scientific) if necessary.

### RNA synthesis and LNP encapsulation

mRNA was produced as previously described^51^. In short, the coding sequence of PE2 and PEmax were cloned into the mRNA production plasmid using HiFi DNA Assembly Master Mix (NEB). mRNAs were transcribed to contain 101 nucleotide-long poly(A) tails. m1Ψ-5′-triphosphate (TriLink) instead of UTP was used to generate modified nucleoside-containing mRNA. Capping of the *in vitro* transcribed mRNAs was performed co-transcriptionally using the trinucleotide cap1 analog, CleanCap (TriLink). mRNA was purified by cellulose (Sigma-Aldrich) purification as described^52^. All mRNAs were analyzed by agarose gel electrophoresis and were stored frozen at −20 °C. Synthetic pegRNAs were ordered and synthesized by Axolabs (peg-mod_1) or Agilent (peg-mod_2-4). LNP were formulated as described previously^53^. In short, an ethanolic solution of 1,2-distearoyl-*sn*-glycero-3-phosphocholine, cholesterol, a PEG lipid and an ionizable cationic lipid was rapidly mixed with an aqueous solution (pH 4) containing prime editor mRNA using an in-line mixer. The lipid and the LNP used in this study are described in patent application WO 2017/004143. The resulting LNP formulation was dialyzed overnight against 1× PBS, 0.2-μm sterile filtered and stored at −80 °C at a concentration of 1 μg/μl of total RNA. Encapsulation efficiencies of mRNA in the LNP were >97 % as measured by the Quant-iT Ribogreen Assay (Life Technologies) and LNP sizes were below 80 nm as measured by a Malvern Zetasizer (Malvern Panalytical).

### Animal studies

Animal experiments were performed in accordance with protocols approved by the Kantonales Veterinäramt Zürich and in compliance with all relevant ethical regulations. *Pah^enu2^* and C57BL/6J mice were housed in a pathogen-free animal facility at the Institute of Pharmacology and Toxicology of the University of Zurich. Mice were kept in a temperature- and humidity-controlled room on a 12-hour light-dark cycle. Mice were fed a standard laboratory chow (Kliba Nafag no. 3437 with 18.5% crude protein) and genotyped at weaning. Heterozygous *Pah^enu2^* littermates were used as controls for physiological L-Phe concentrations in the blood (<120 μmol/L). For sampling of blood for L-Phe determination, mice were fasted for 3 to 4 hours, and the blood was collected from the tail vein. Adult mice were injected with 5-10 × 10^13^ vg/kg (AAV) or with 1 – 3 mg/kg (LNP) in a maximal volume of 150 µl via the tail vein. The selected AAV and LNP doses were based on the maximum injection volume for adults (150 μl of undiluted viral vectors via the tail vein).

### Primary hepatocyte isolation

Primary hepatocytes were isolated using a two-step perfusion method. Briefly, pre-perfusion with Hanks’ buffer (supplemented with 0.5 mM EDTA and 25 mM Hepes) was performed by inserting the cannula through the superior vena cava and cutting the portal vein. Next, livers were perfused at low flow for about 10 min with digestion buffer (low-glucose DMEM supplemented with 1 mM Hepes) containing freshly added Liberase (32 μg/ml; Roche). Digestion was stopped using isolation buffer (low-glucose DMEM supplemented with 10% FBS), and cells were separated from the matrix by gently pushing with a cell scraper. The cell suspension was filtered through a 100-μm filter (Corning), and hepatocytes were purified by two low-speed centrifugation steps (50g for 2 min).

### PCR amplification for deep sequencing

Genomic DNA from mouse tissues were isolated by direct lysis. Locus-specific primers were used to generate targeted amplicons for deep sequencing. First, input genomic DNA was amplified in a 10-μl reaction for 26 cycles using NEBNext High-Fidelity 2x PCR Master Mix (NEB). PCR products were purified using Sera-Mag magnetic beads (Cytiva) and subsequently amplified for six cycles using primers with sequencing adapters. Approximately equal amounts of PCR products from each sample were pooled, gel purified, and quantified using a Qubit 3.0 fluorometer and the dsDNA HS Assay Kit (Thermo Fisher Scientific). Paired-end sequencing of purified libraries was performed on an Illumina MiSeq.

### NGS data analysis

Sequencing reads were demultiplexed using MiSeq Reporter (Illumina). Amplicon sequences were aligned to their reference sequences using CRISPResso2^54^. Prime editing efficiencies were calculated as percentage of (number of reads containing only the desired edit)/(number of total reads). Indel yields were calculated as percentage of (number of indel-containing reads)/(total reads).

### Quantification of phenylalanine in the blood

Amino acids were extracted from a 3.2-mm dried blood sample using the Neomass AAAC Plus newborn screening kit (Labsystems Diagnostics). A UHPLC (ultrahigh performance liquid chromatography) Nexera X2 coupled to an LCMS-8060 triple quadrupole mass spectrometer with electrospray ionization (Shimadzu) was used for the quantitative analysis of phenylalanine. LabSolutions and Neonatal Solution software (Shimadzu) were used for data acquisition and data analysis.

### Quantification of phenylalanine enzyme activity

Whole liver extracts were analyzed using isotope-dilution liquid chromatography-electrospray ionization tandem mass spectrometry (LC-ESI-MS/MS) as described previously^55^.

### Western blot

Proteins were isolated from liver samples of treated and untreated animals. Briefly, cells were lysed in RIPA buffer, supplemented with protease inhibitors (Sigma-Aldrich). Protein amounts were determined using the Pierce BCA Protein Assay Kit (Thermo Fisher). Equal amounts of protein (80 μg) were separated by SDS-PAGE (Thermo Fisher) and transferred to a 0.45 μm nitrocellulose membrane (Amersham). Membranes were incubated with mouse anti-Cas9 (1:1’000; Cat. No. #14697T; Cell Signaling) and rabbit anti-GAPDH (1:10’000; Cat. No. ab181602; abcam). Signals were detected by fluorescence using IRDye-conjugated secondary antibodies (Licor).

### RNA isolation and RT-qPCR

RNA was isolated from shock frozen liver samples using the RNeasy Mini kit (Qiagen) according to the manufacturer’s instructions. RNA was reverse transcribed to cDNA using random primers and GoScript Reverse Transcriptase kit (Promega). RT-qPCR was performed using Firepol qPCR Master Mix (Solis BioDyne) and analyzed by 7900HT Fast Real-Time PCR System (Applied Biosystems). Fold changes were calculated using the delta Ct method. Used primers are listed in Table S3.

### Immunofluorescence

During liver perfusion, one liver lobe was tightened off using a silk suture thread (Fine Science Tools, FST). Tissues were transferred to a 30% sucrose solution overnight at 4 °C and embedded in OCT compound in cryomolds (Tissue-Tek) and frozen at -80°C for at least 30 minutes. Frozen tissues were sectioned at 7 µm at −20 °C, and mounted directly on Superfrost Plus slides (Thermo Fisher Scientific). Cryosections were counterstained with DAPI (Thermo Fisher Scientific) and mounted in VECTASHIELD mounting medium (Vector Labs). Two frozen sections were analyzed per mouse per tissue. Mouse tissue was imaged using Zeiss Axioscope and Colibri 7 LED Illumination lighting system. Imaging conditions and intensity scales were matched for all images. Images were taken using Zeiss software Zen2 and analyzed by Fiji ImageJ software (v1.51n)^56^.

### Histology

Livers were fixed using 4% paraformaldehyde (PFA, Sigma-Aldrich), followed by ethanol dehydration and paraffinization. Paraffin blocks were cut into 5 μm thick sections, deparaffinized with xylene, and rehydrated. Sections were HE-stained and examined for histopathological changes.

### Detection of plasma pro-inflammatory and damage markers

Blood was collected from the inferior vena cava using Lithium-Heparin coated 0.5 ml tubes (MiniCollect) prior to liver perfusion. Samples were centrifuged at 2000xg for 10 min and the supernatant was collected and stored at -20°C until measurement. AST and ALT levels from all mouse samples were measured as routine parameters at the Division of Clinical Chemistry and Biochemistry at the University Children’s Hospital Zurich using Alinity ci-series. Pro-inflammatory cytokines were detected using LEGENDplex Mouse Inflammation panel (13-plex; Biolgend; catalog number 740446; lot number B354399), a bead-based multiplex assay, according to the manufacturer’s instructions.

### Statistical analysis

Statistical analyses were performed using GraphPad Prism 9.0.0 for macOS. Data are represented as biological replicates and are depicted as means ± standard deviation (s.d.) or standard error of the mean (s.e.m.) as indicated in the corresponding figure legends. Likewise, sample sizes and the statistical tests used are described in detail in the respective figure legends. For all analyses, P < 0.05 was considered statistically significant.

## Acknowledgments

We thank the Functional Genomics Center Zurich for technical support and access to instruments at the University of Zurich and ETH Zürich, and members of the Thöny, Häberle and Schwank labs for discussions. We thank the viral vector facility (VVF) Zurich for scientific advice and providing plasmids containing ssAAV2 and scAAV2 genomes. pU6-tevopreq1-GG-acceptor (Addgene No.174038) and pCMV-PEmax (Addgene No. 174820) were gifts from D. Liu and AAV capsid and helper plasmids (Addgene No. 112865 and 112867) were gifts from J.M. Wilson. We thank Daniel Ryan and the whole Agilent team for providing PACE-modified synthetic pegRNAs. This study was supported by the Swiss National Science Foundation (SNSF) grant no. 310030_185293 (to G.S.), the SERI funded ERC-CoG ‘GeneRepair’ (to G.S), SNSF Sinergia grant no. 180257 (to B.T.), Novartis Foundation for Medical-Biological Research no. FN20-0000000203 (to D.B.), SNSF Spark fellowship no. 196287 (to D.B.), the University Research Priority Programs ‘ITINERARE’ (to D.B and G.S) and ‘Human Reproduction Reloaded’ (G.S. and E.I), the Promedica Foundation (to G.S.) and the University Zürich Candoc grant no. FK-22-033 (to T.R.).

## Author contributions

T.R. and G.S. designed the study. T.R., E.I, A.T. and L.V. performed and/or analyzed *in vivo* experiments. T.R., Y.W., D.B., E.V. and E.I. performed and analyzed *in vitro* experiments. N.R. and E.F. performed blood Phe and *Pah* measurements. C.S. performed genotyping experiments. L.S., K.F.M., T.R., Y.W. and D.B. performed molecular cloning experiments. D.B performed western blotting analysis. N.P., H.M. and M.V. performed *in vitro* transcription of mRNA. Y.K.T., P.J.C.L., J.M. and S.H.Y.F. developed LNP formulations and complexed mRNAs with LNP. M.M performed measurements for toxicity and proinflammatory markers. A.C. performed AST and ALT measurements. B.T., M.K. and J.H. provided technical and conceptual advice. T.R. prepared figures. T.R. and G.S. wrote the manuscript. All authors reviewed the manuscript.

## Data availability

All data associated with this study are present in the paper or the Supplementary Materials. Illumina sequencing data are available in the Sequence Read Archive (SRA) under the accession number PRJNA947564.

## Supplementary Material

**Figure S1)** *In vitro* optimization of PE components for correction of the disease-causing T-to-C mutation at the *Pah^enu^*locus.

**Figure S2)** *In vivo* prime editing using AAV-mediated PE delivery in PKU mice.

**Figure S3)** Expression kinetics of the PE after LNP-mRNA delivery into the liver.

**Figure S4)** *In vivo* prime editing rates in mice treated with AAV-pegRNA and LNP-mRNA.

**Figure S5)** Editing rates in non-liver tissues and liver histology at different timepoints after LNP-mRNA delivery.

**Figure S6)** Gating strategy identifying proinflammatory markers.

**Note 1)** Amino acid sequences of AAV vectors used in this study.

**Note 2)** Complete image of Western blot.

**Table 1)** pegRNA designs tested for correction of the *Pah^enu2^* disease locus.

**Table 2)** Oligonucleotides used for this study.

**Table 3)** Oligonucleotides used for RT-qPCR.

**Table 4)** modified pegRNAs used in this study.

