## Supplementary figures for "Treatment of a genetic liver disease in mice through transient prime editor expression"

**Supplementary Material**

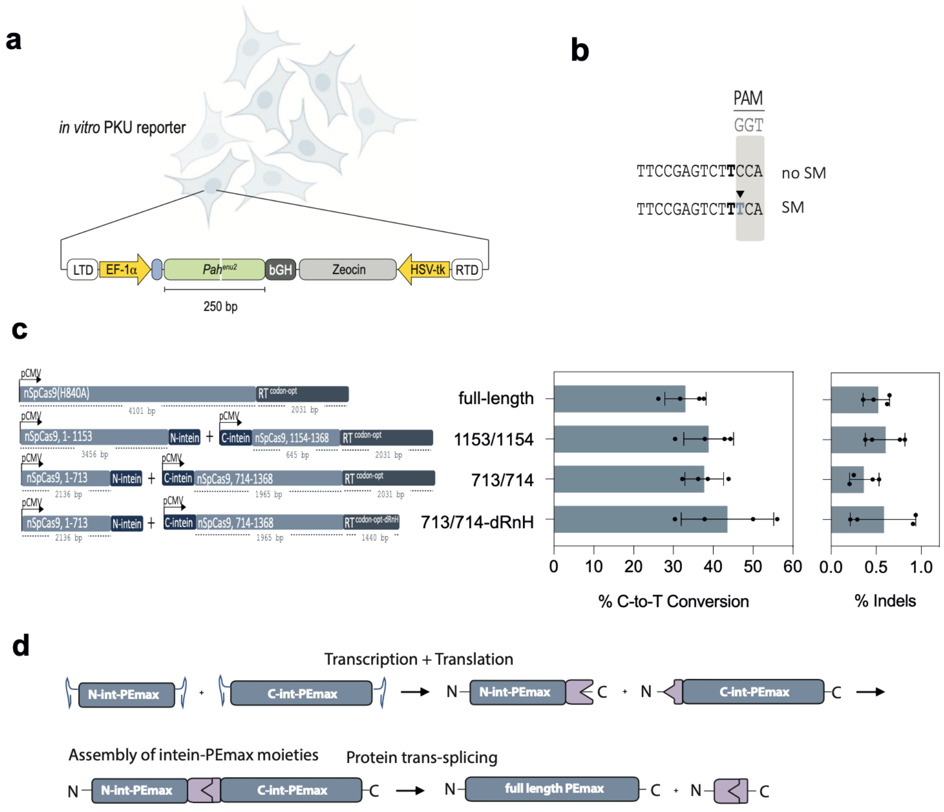


**Figure S1 | *In vitro* optimization of PE components for correction of the disease-causing T-to-C mutation at the *Pah$^{enu}$* locus.** (**a**) Schematic representation of HEK293T or K562 Pah$^{enu2}$ reporter cell lines, generated using the PiggyBac transposon system. The pegRNA binding site is indicated by a white line. (**b**) Schematic representation of the RTT in the pegRNAs mPKU-2.1 (noSM) or mPKU-SM (SM). Bold letter indicates the base that corrects the pathogenic mutation. (**c**) Schematic representation of intein-split PEmax constructs (left panel) and editing rates with different variants tested in HEK293T cells with the integrated Pah$^{enu2}$ locus. Percentage of intended C-to-T conversions (middle panel) and indels (right panel). (**d**) Schematic of the intein-split trans-splicing system. dRnH: PEmax without the RnaseH domain in the RT. RTD, right and LTD, left terminal domains; EF-1a, Human elongation factor-1 alpha promoter, *Pah*, Phenylalanine hydroxylase; bGH, bovine growth hormone polyadenylation signal; HSV-tk, herpes simplex virus thymidine kinase promoter; elements are not depicted to scale. Values represent mean +/- s.d. of four independent biological replicates. Means were compared using an unpaired Student's t tests. (ns, not significant, P > 0.05).

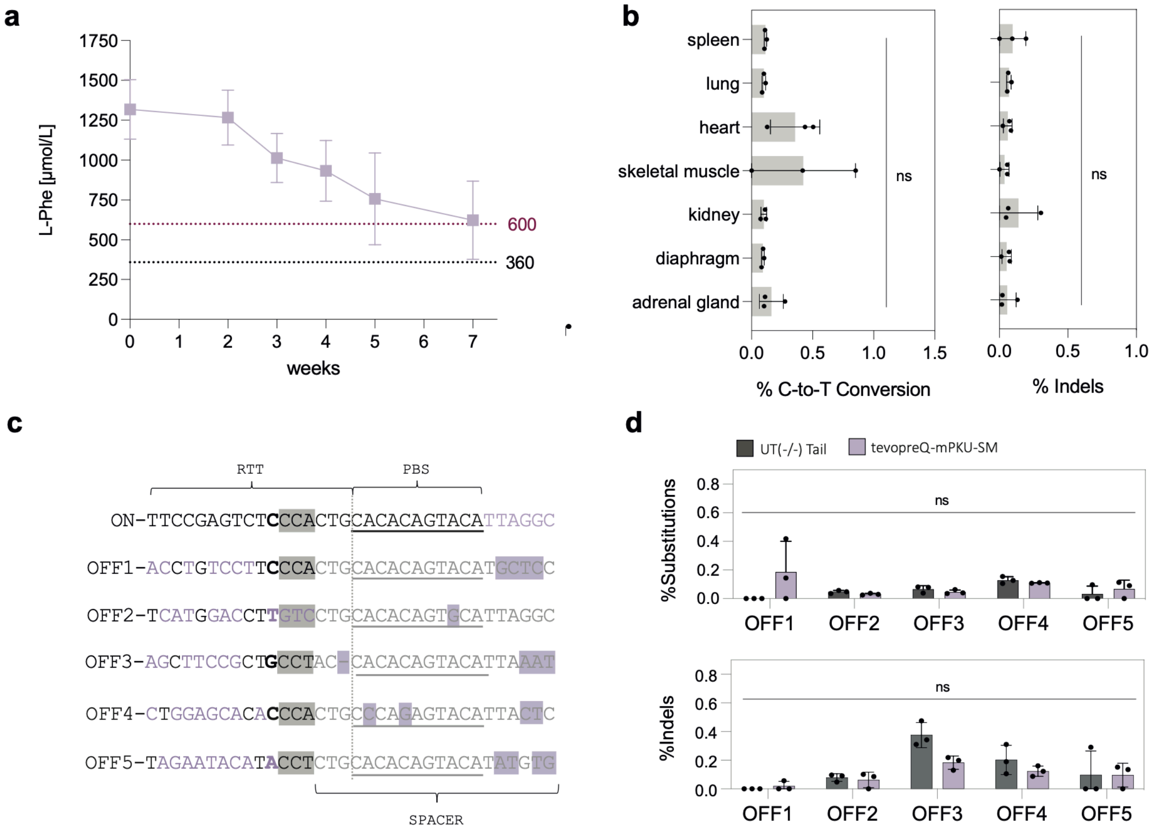


**Figure S2 | *In vivo* prime editing using AAV-mediated PE delivery in PKU mice.**

(**a**) Change of Phe levels over time in animals treated with AAVs encoding for 713/714 Pemax-dRnH and tevopreQ$_1$-mPKU-SM (n=3). Values represent mean +/- SEM of independent biological replicates. (**b**) C-to-T conversion rates (left panel) and indel rates (right panel) at the Pah$^{enu2}$ locus in tissues other than the liver from animals treated with 1x10$^{14}$ vg/kg AAV encoding for Pemax. Editing rates were assessed by targeted amplicon sequencing of whole tissue lysates. Means were compared using an ordinary one-way ANOVA using Šídák's multiple comparisons test. (**c-d**) Targeted amplicon sequencing of the top five off-target sites of the mPKU-SM pegRNA identified by CHANGE-seq in Böck et al.[19] (**c**) Indicated are the spacer, the primer binding site (underlined, PBS) and reverse transcriptase template (RTT) of the target site (ON) and 5 off-target sites (OFF1-OFF5). Mismatches to the target site are highlighted in purple. The bases complementary to the PAM site are highlighted in grey. The site of the DNA nick is indicated by a dotted line. (**d**) Indel rates and substitution rates were quantified for tail tissue (UT (-/-) Tail) and isolated hepatocytes from animals treated with dual-AAV with tevopreQ1-mPKU-SM. Means were compared using an ordinary one-way ANOVA using Šídák's multiple comparisons test. Unless otherwise states, values depict mean +/- s.d. of independent biological replicates (ns, not significant, P > 0.05).

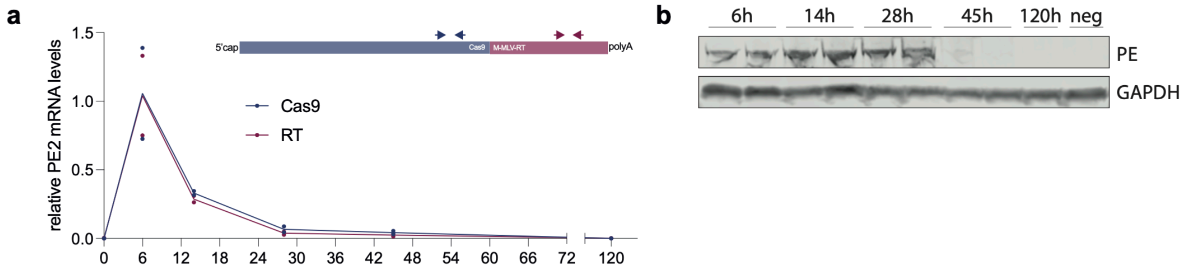


**Figure S3 | Expression kinetics of the PE after LNP-mRNA delivery into the liver.** (**a**) Expression kinetics of PE mRNA delivered via LNP at a dose of 2 mg/kg. Relative values were normalized to the housekeeping gene RPLP0 and the average observed peak expression at 6 h.p.i. Each value represents mean of two individual biological replicates. The prime editor transcript was quantified separately using two targeting primers, where one binds the Cas9 (blue) and the other the RT (magenta) domain of the prime editor transcript. (**b**) Expression kinetics of the PE protein. Relative values were normalized to the housekeeping gene GAPDH and the average observed peak expression at 28 h.p.i. Each value represents mean of two individual biological replicates. The full western plot image is shown in supplementary note 2.

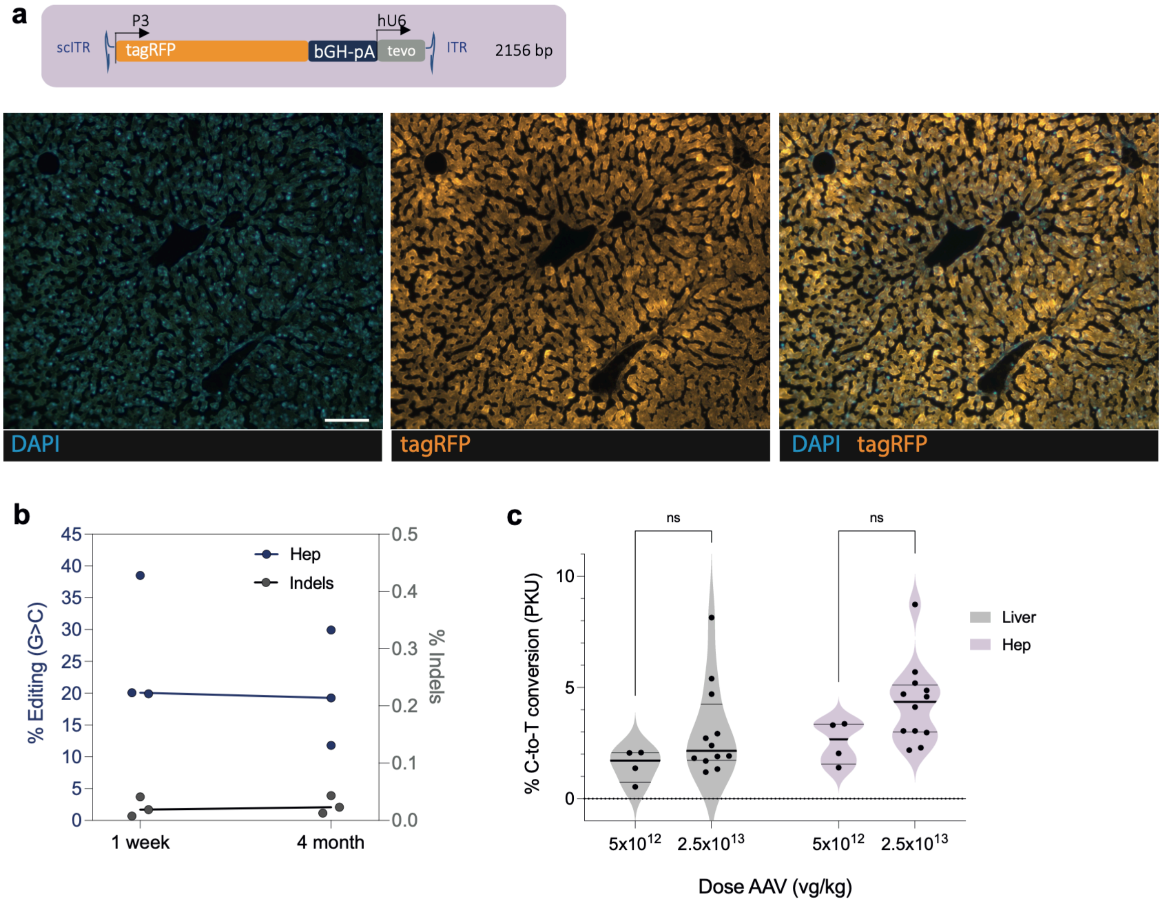


**Figure S4 | *In vivo* prime editing rates in mice treated with AAV-pegRNA and LNP-mRNA.** (**a**) Illustration of the self-complementary (sc) AAV genome expressing tevopreQ1-modified pegRNAs. ITR: inverted terminal repeat; P3: liver-specific P3 promoter; tagRFP: tag red fluorescent protein; bGH-pA: bovine growth hormone polyadenylation signal, hU6: human U6 promoter, tevo: trimmed evopreQ$_1$-modified pegRNA (epegRNA). tagRFP expressing cells in the mouse liver 3 weeks after injection of scAAV at a dose of 2.5x$10^{13}$ vg/kg. Scale bar: 50 µm. (**b**) Editing rates at the *Dnmt1* locus 1 week and 4 months after LNP-mRNA injection. Animals were pre-treated with the same scAAV-pegRNA dose. Lines connect medians and dots represent individual animals. Correction rates are shown in dark blue and one the left y-axis. Indel rates are shown in grey and the right y-axis. (**c**) Comparison of C-to-T conversion rates at the *Pah$^{enu2}$* locus using pretreatment with either 5x$10^{12}$ or 2.5x$10^{13}$ vg/kg scAAV expressing tevopreQ$_1$-mPKU-SM with subsequent three-times redosing of LNP expressing PEmax. Individual data points and medians are depicted from whole liver isolates and primary isolated hepatocytes of the same animals. Values were compared using Šídák's multiple comparisons test. ns, not significant.

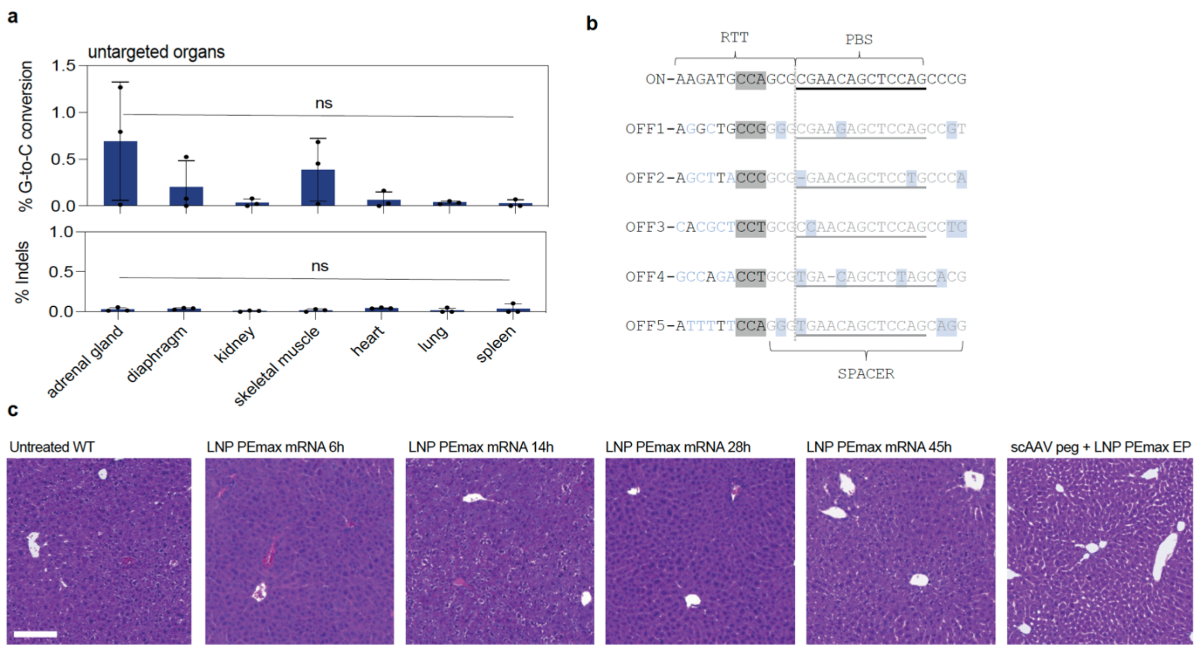


**Figure S5 | Editing rates in non-liver tissues and liver histology at different timepoints after LNP-mRNA delivery.** (**a**) C-to-T conversion rates (upper panel) and indel rates (lower panel) at the *Dnmt1* locus on genomic DNA isolated from tissues other than the liver from (n=3) animals pretreated with scAAV encoding for the tevopreQ$_1$-modified pegRNA targeting *Dnmt1* and dosed once with 3 mg/kg PE2 or PEmax mRNA. Editing rates were assessed by NGS. (**b**) Top five off-target sites of the *Dnmt1*-targeting tevopreQ$_1$-modified pegRNA identified by CHANGE-seq[16]. Indicated are the spacer, the primer binding site (underlined, PBS) and reverse transcriptase template (RTT) of the target site (ON) and off-target sites (OFF1-OFF5). Mismatches to the target site are highlighted in light blue. The bases complimentary to the PAM site are highlighted in grey. The site of the DNA nick is indicated by a dotted line. (**c**) Histological images of hematoxylin and eosin (H&E) stained 5µm thick liver tissue. Images are representative of two individual animals. EP: Experimental endpoint of scAAVpeg + LNP PEmax is one week after LNP delivery. Scale bar: 100 µm.

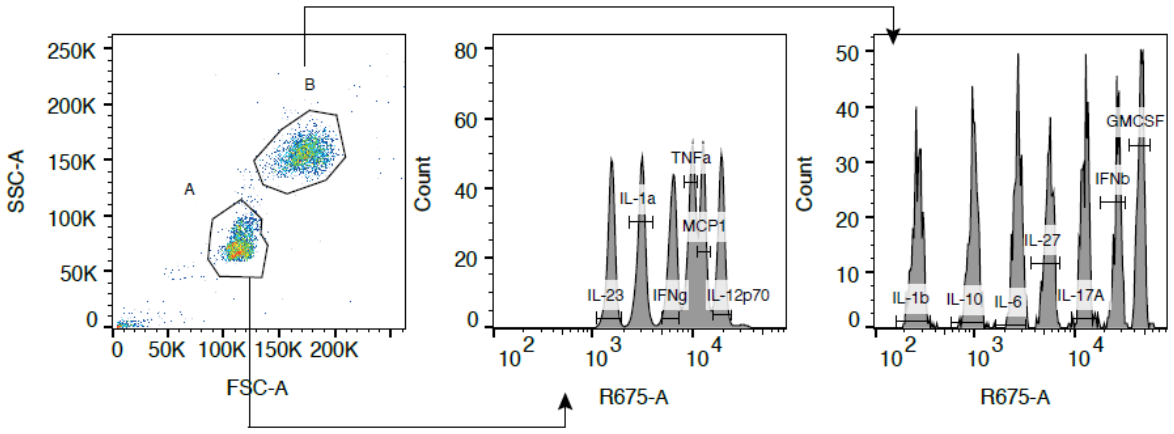


**Figure S6 | Gating strategy identifying proinflammatory markers.**

**Supplementary Note 1 |** Amino acid sequences of AAV vectors used in this study.

| |
|---|
| **N-int-PEmax (1-712):** BPSV40NLS – Cas9 (1-712) – (GGGGS)3 linker – Npu N-intein – SV40-NLS |
| MKRTADGSEFESPKKKRKVDKKYSIGLDIGTNSVGWAVITDEYKVPSKKFKVLGNTDRHSIKKNLIGALLFDSG<br>ETAEATRLKRTARRRYTRRKNRICYLQEIFSNEMAKVDDSFFHRLEESFLVEEDKKHERHPIFGNIVDEVAYHEK<br>YPTIYHLRKKLVDSTDKADLRLIYLALAHMIKFRGHFLIEGDLNPDNSDVDKLFIQLVQTYNQLFEENPINASGV<br>DAKAILSARLSKSRKLENLIAQLPGEKKNGLFGNLIALSLGLTPNFKSNFDLAEDAKLQLSKDTYDDDLDNLLAQ<br>IGDQYADLFLAAKNLSDAILLSDILRVNTEITKAPLSASMIKRYDEHHQDLTLLKALVRQQLPEKYKEIFFDQSKN<br>GYAGYIDGGASQEEFYKFIKPILEKMDGTEELLVKLKREDLLRKQRTFDNGSIPHQIHLGELHAILRRQEDFYPFL<br>KDNREKIEKILTFRIPYYVGPLARGNSRFAWMTRKSEETITPWNFEEVVDKGASAQSFIERMTNFDKNLPNEKVL<br>PKHSLLYEYFTVYNELTKVKYVTEGMRKPAFLSGEQKKAIVDLLFKTNRKVTVKQLKEDYFKKIECFDSVEISG<br>VEDRFNASLGTYHDLLKIIKDKDFLDNEENEDILEDIVLTLTLFEDREMIEERLKTYAHLFDDKVMKQLKRRRYT<br>GWGRLSRKLINGIRDKQSGKTILDFLKSDGFANRNFMQLIHDDSLTFKEDIQKAQVGGGGSGGGGSGGGGSCLS<br>YETEILTVEYGLLPIGKIVEKRIECTVYSVDNNGNIYTQPVAQWHDRGEQEVFEYCLEDGSLIRATKDHKFMTVD<br>GQMLPIDEIFERELDLMRVDNLPNEFEPKKKRKV* |
| **C-int-PEmax (713-1368):** BPSV40NLS – Npu C-intein – (GGGGS)3 linker – Cas9 (713-1368) – (SGGS)2 –<br>BPSV40NLS – (SGGS)2 – M-MLV-RT-dRnH – SGGS – BPSV40NLS – cMyc NLS |
| MKRTADGSEFESPKKKRKVIKIATRKYLGKQNVYDIGVERDHNFALKNGFIASNSGGGGSGGGGSGGGGSSGQ<br>GDSLHEHIANLAGSPAIKKGILQTVKVVDELVKVMGRHKPENIVIEMARENQTTQKGQKNSRERMKRIEEGIKE<br>LGSQILKEHPVENTQLQNEKLYLYYLQNGRDMYVDQELDINRLSDYDVDAIVPQSFLKDDSIDNKVLTRSDKNR<br>GKSDNVPSEEVVKKMKNYWRQLLNAKLITQRKFDNLTKAERGGLSELDKAGFIKRQLVETRQITKHVAQILDSR<br>MNTKYDENDKLIREVKVITLKSKLVSDFRKDFQFYKVREINNYHHAHDAYLNAVVGTALIKKYPKLESEFVYG<br>DYKVYDVRKMIAKSEQEIGKATAKYFFYSNIMNFFKTEITLANGEIRKRPLIETNGETGEIVWDKGRDFATVRKV<br>LSMPQVNIVKKTEVQTGGFSKESILPKRNSDKLIARKKDWDPKKYGGFDSPTVAYSVLVVAKVEKGKSKKLKS<br>VKELLGITIMERSSFEKNPIDFLEAKGYKEVKKDLIIKLPKYSLFELENGRKRMLASAGELQKGNELALPSKYVNF<br>LYLASHYEKLKGSPEDNEQKQLFVEQHKHYLDEIIEQISEFSKRVILADANLDKVLSAYNKHRDKPIREQAENIIH<br>LFTLTNLGAPAAFKYFDTTIDRKRYTSTKEVLDATLIHQSITGLYETRIDLSQLGGDSGGSSGGSKRTADGSEFES<br>PKKKRKVSGGSSGGSTLNIEDEYRLHETSKEPDVSLGSTWLSDFPQAWAETGGMGLAVRQAPLIIPLKATSTPVS<br>IKQYPMSQEARLGIKPHIQRLLDQGILVPCQSPWNTPLLPVKKPGTNDYRPVQDLREVNKRVEDIHPTVPNPYNL<br>LSGLPPSHQWYTVLDLKDAFFCLRLHPTSQPLFAFEWRDPEMGISGQLTWTRLPQGFKNSPTLFNEALHRDLAD<br>FRIQHPDLILLQYVDDLLLAATSELDCQQGTRALLQTLGNLGYRASAKKAQICQKQVKYLGYLLKEGQRWLTE<br>ARKETVMGQPTPKTPRQLREFLGKAGFCRLFIPGFAEMAAPLYPLTKPGTLFNWGPDQQKAYQEIKQALLTAPA<br>LGLPDLTKPFELFVDEKQGYAKGVLTQKLGPWRRPVAYLSKKLDPVAAGWPPCLRMVAAIAVLTKDAGKLTM<br>GQPLVILAPHAVEALVKQPPDRWLSNARMTHYQALLLDTDRVQFGPVVALNPSGGSKRTADGSEFESPKKKRK<br>VGSGPAAKRVKLD* |
| **N-int-PEmax (1-1153):** BPSV40NLS – Cas9 (1-712) – (GGGGS)3 linker – Npu N-intein – SV40-NLS |
| MKRTADGSEFESPKKKRKVDKKYSIGLDIGTNSVGWAVITDEYKVPSKKFKVLGNTDRHSIKKNLIGALLFDSG<br>ETAEATRLKRTARRRYTRRKNRICYLQEIFSNEMAKVDDSFFHRLEESFLVEEDKKHERHPIFGNIVDEVAYHEK<br>YPTIYHLRKKLVDSTDKADLRLIYLALAHMIKFRGHFLIEGDLNPDNSDVDKLFIQLVQTYNQLFEENPINASGV<br>DAKAILSARLSKSRKLENLIAQLPGEKKNGLFGNLIALSLGLTPNFKSNFDLAEDAKLQLSKDTYDDDLDNLLAQ<br>IGDQYADLFLAAKNLSDAILLSDILRVNTEITKAPLSASMIKRYDEHHQDLTLLKALVRQQLPEKYKEIFFDQSKN<br>GYAGYIDGGASQEEFYKFIKPILEKMDGTEELLVKLKREDLLRKQRTFDNGSIPHQIHLGELHAILRRQEDFYPFL<br>KDNREKIEKILTFRIPYYVGPLARGNSRFAWMTRKSEETITPWNFEEVVDKGASAQSFIERMTNFDKNLPNEKVL<br>PKHSLLYEYFTVYNELTKVKYVTEGMRKPAFLSGEQKKAIVDLLFKTNRKVTVKQLKEDYFKKIECFDSVEISG<br>VEDRFNASLGTYHDLLKIIKDKDFLDNEENEDILEDIVLTLTLFEDREMIEERLKTYAHLFDDKVMKQLKRRRYT<br>GWGRLSRKLINGIRDKQSGKTILDFLKSDGFANRNFMQLIHDDSLTFKEDIQKAQVSGQGDSLHEHIANLAGSPA<br>IKKGILQTVKVVDELVKVMGRHKPENIVIEMARENQTTQKGQKNSRERMKRIEEGIKELGSQILKEHPVENTQLQ<br>NEKLYLYYLQNGRDMYVDQELDINRLSDYDVDAIVPQSFLKDDSIDNKVLTRSDKNRGKSDNVPSEEVVKKMK<br>NYWRQLLNAKLITQRKFDNLTKAERGGLSELDKAGFIKRQLVETRQITKHVAQILDSRMNTKYDENDKLIREVK<br>VITLKSKLVSDFRKDFQFYKVREINNYHHAHDAYLNAVVGTALIKKYPKLESEFVYGDYKVYDVRKMIAKSEQ<br>EIGKATAKYFFYSNIMNFFKTEITLANGEIRKRPLIETNGETGEIVWDKGRDFATVRKVLSMPQVNIVKKTEVQT<br>GGFSKESILPKRNSDKLIARKKDWDPKKYGGFDSPTVAYSVLVVAKVEKGKGGGGSGGGGSGGGGSCLSYETE<br>ILTVEYGLLPIGKIVEKRIECTVYSVDNNGNIYTQPVAQWHDRGEQEVFEYCLEDGSLIRATKDHKFMTVDGQM<br>LPIDEIFERELDLMRVDNLPN* |
| **C-int-PEmax (1154-1368):** BPSV40NLS – Npu C-intein – (GGGGS)3 linker – Cas9 (713-1368) – (SGGS)2 –<br>BPSV40NLS – (SGGS)2 – M-MLV-RT-dRnH – SGGS – BPSV40NLS – cMyc NLS |
| MKRTADGSEFESPKKKRKVIKIATRKYLGKQNVYDIGVERDHNFALKNGFIASNSGGGGSGGGGSGGGGSSKK<br>LKSVKELLGITIMERSSFEKNPIDFLEAKGYKEVKKDLIIKLPKYSLFELENGRKRMLASAGELQKGNELALPSKY |

| |
|---|
| VNFLYLASHYEKLKGSPEDNEQKQLFVEQHKHYLDEIIEQISEFSKRVILADANLDKVLSAYNKHRDKPIREQAE<br>NIIHLFTLTNLGAPAAFKYFDTTIDRKRYTSTKEVLDATLIHQSITGLYETRIDLSQLGGDSGGSSGGSKRTADGSE<br>FESPKKKRKVSGGSSGGSTLNIEDEYRLHETSKEPDVSLGSTWLSDFPQAWAETGGMGLAVRQAPLIIPLKATST<br>PVSIKQYPMSQEARLGIKPHIQRLLDQGILVPCQSPWNTPLLPVKKPGTNDYRPVQDLREVNKRVEDIHPTVPNP<br>YNLLSGLPPSHQWYTVLDLKDAFFCLRLHPTSQPLFAFEWRDPEMGISGQLTWTRLPQGFKNSPTLFNEALHRD<br>LADFRIQHPDLILLQYVDDLLLAATSELDCQQGTRALLQTLGNLGYRASAKKAQICQKQVKYLGYLLKEGQRW<br>LTEARKETVMGQPTPKTPRQLREFLGKAGFCRLFIPGFAEMAAPLYPLTKPGTLFNWGPDQQKAYQEIKQALLT<br>APALGLPDLTKPFELFVDEKQGYAKGVLTQKLGPWRRPVAYLSKKLDPVAAGWPPCLRMVAAIAVLTKDAGK<br>LTMGQPLVILAPHAVEALVKQPPDRWLSNARMTHYQALLLDTDRVQFGPVVALNPSGGSKRTADGSEFESPKK<br>KRKVGSGPAAKRVKLD* |
| **tagRFP** |
| MVSKGEELIKENMHMKLYMEGTVNNHHFKCTSEGEGKPYEGTQTMRIKVVEGGPLPFAFDILATSFMYGSRTFI<br>NHTQGIPDFFKQSFPEGFTWERVTTYEDGGVLTATQDTSLQDGCLIYNVKIRGVNFPSNGPVMQKKTLGWEANT<br>EMLYPADGGLEGRSDMALKLVGGGHLICNFKTTYRSKKPAKNLKMPGVYYVDHRLERIKEADKETYVEQHEV<br>AVARYCDLPSKLGHKLN* |
| **PE2 mRNA production vector (full length PE2):** BPSV40NLS – Cas9 (H840A; PE2) – (SGGS)2 – XTEN – (SGGS)2 – M-MLV-RT (PE2) – SGGS – BPSV40NLS |
| MKRTADGSEFESPKKKRKVDKKYSIGLDIGTNSVGWAVITDEYKVPSKKFKVLGNTDRHSIKKNLIGALLFDSG<br>ETAEATRLKRTARRRYTRRKNRICYLQEIFSNEMAKVDDSFFHRLEESFLVEEDKKHERHPIFGNIVDEVAYHEK<br>YPTIYHLRKKLVDSTDKADLRLIYLALAHMIKFRGHFLIEGDLNPDNSDVDKLFIQLVQTYNQLFEENPINASGV<br>DAKAILSARLSKSRRLENLIAQLPGEKKNGLFGNLIALSLGLTPNFKSNFDLAEDAKLQLSKDTYDDDLDNLLAQ<br>IGDQYADLFLAAKNLSDAILLSDILRVNTEITKAPLSASMIKRYDEHHQDLTLLKALVRQQLPEKYKEIFFDQSKN<br>GYAGYIDGGASQEEFYKFIKPILEKMDGTEELLVKLNREDLLRKQRTFDNGSIPHQIHLGELHAILRRQEDFYPFL<br>KDNREKIEKILTFRIPYYVGPLARGNSRFAWMTRKSEETITPWNFEEVVDKGASAQSFIERMTNFDKNLPNEKVL<br>PKHSLLYEYFTVYNELTKVKYVTEGMRKPAFLSGEQKKAIVDLLFKTNRKVTVKQLKEDYFKKIECFDSVEISG<br>VEDRFNASLGTYHDLLKIIKDKDFLDNEENEDILEDIVLTLTLFEDREMIEERLKTYAHLFDDKVMKQLKRRRYT<br>GWGRLSRKLINGIRDKQSGKTILDFLKSDGFANRNFMQLIHDDSLTFKEDIQKAQVSGQGDSLHEHIANLAGSPA<br>IKKGILQTVKVVDELVKVMGRHKPENIVIEMARENQTTQKGQKNSRERMKRIEEGIKELGSQILKEHPVENTQLQ<br>NEKLYLYYLQNGRDMYVDQELDINRLSDYDVDAIVPQSFLKDDSIDNKVLTRSDKNRGKSDNVPSEEVVKKMK<br>NYWRQLLNAKLITQRKFDNLTKAERGGLSELDKAGFIKRQLVETRQITKHVAQILDSRMNTKYDENDKLIREVK<br>VITLKSKLVSDFRKDFQFYKVREINNYHHAHDAYLNAVVGTALIKKYPKLESEFVYGDYKVYDVRKMIAKSEQ<br>EIGKATAKYFFYSNIMNFFKTEITLANGEIRKRPLIETNGETGEIVWDKGRDFATVRKVLSMPQVNIVKKTEVQT<br>GGFSKESILPKRNSDKLIARKKDWDPKKYGGFDSPTVAYSVLVVAKVEKGKSKKLKSVKELLGITIMERSSFEK<br>NPIDFLEAKGYKEVKKDLIIKLPKYSLFELENGRKRMLASAGELQKGNELALPSKYVNFLYLASHYEKLKGSPED<br>NEQKQLFVEQHKHYLDEIIEQISEFSKRVILADANLDKVLSAYNKHRDKPIREQAENIIHLFTLTNLGAPAAFKYF<br>DTTIDRKRYTSTKEVLDATLIHQSITGLYETRIDLSQLGGDSGGSSGGSSGSETPGTSESATPESSGGSSGGSSTLNI<br>EDEYRLHETSKEPDVSLGSTWLSDFPQAWAETGGMGLAVRQAPLIIPLKATSTPVSIKQYPMSQEARLGIKPHIQ<br>RLLDQGILVPCQSPWNTPLLPVKKPGTNDYRPVQDLREVNKRVEDIHPTVPNPYNLLSGLPPSHQWYTVLDLKD<br>AFFCLRLHPTSQPLFAFEWRDPEMGISGQLTWTRLPQGFKNSPTLFNEALHRDLADFRIQHPDLILLQYVDDLLL<br>AATSELDCQQGTRALLQTLGNLGYRASAKKAQICQKQVKYLGYLLKEGQRWLTEARKETVMGQPTPKTPRQL<br>REFLGKAGFCRLFIPGFAEMAAPLYPLTKPGTLFNWGPDQQKAYQEIKQALLTAPALGLPDLTKPFELFVDEKQ<br>GYAKGVLTQKLGPWRRPVAYLSKKLDPVAAGWPPCLRMVAAIAVLTKDAGKLTMGQPLVILAPHAVEALVKQ<br>PPDRWLSNARMTHYQALLLDTDRVQFGPVVALNPATLLPLPEEGLQHNCLDILAEAHGTRPDLTDQPLPDADHT<br>WYTDGSSLLQEGQRKAGAAVTTETEVIWAKALPAGTSAQRAELIALTQALKMAEGKKLNVYTDSRYAFATAHI<br>HGEIYRRRGWLTSEGKEIKNKDEILALLKALFLPKRLSIIHCPGHQKGHSAEARGNRMADQAARKAAITETPDTS<br>TLLIENSSPSGGSKRTADGSEFEPKKKRKV* |
| **PE2 mRNA production vector (full length PEmax):** BPSV40NLS – Cas9 (H840A; PEmax) – (SGGS)2 – BPSV40NLS – (SGGS)2 – M-MLV-RT (PEmax) – SGGS – BPSV40NLS – cMyc NLS |
| MKRTADGSEFESPKKKRKVDKKYSIGLDIGTNSVGWAVITDEYKVPSKKFKVLGNTDRHSIKKNLIGALLFDSG<br>ETAEATRLKRTARRRYTRRKNRICYLQEIFSNEMAKVDDSFFHRLEESFLVEEDKKHERHPIFGNIVDEVAYHEK<br>YPTIYHLRKKLVDSTDKADLRLIYLALAHMIKFRGHFLIEGDLNPDNSDVDKLFIQLVQTYNQLFEENPINASGV<br>DAKAILSARLSKSRKLENLIAQLPGEKKNGLFGNLIALSLGLTPNFKSNFDLAEDAKLQLSKDTYDDDLDNLLAQ<br>IGDQYADLFLAAKNLSDAILLSDILRVNTEITKAPLSASMIKRYDEHHQDLTLLKALVRQQLPEKYKEIFFDQSKN<br>GYAGYIDGGASQEEFYKFIKPILEKMDGTEELLVKLKREDLLRKQRTFDNGSIPHQIHLGELHAILRRQEDFYPFL<br>KDNREKIEKILTFRIPYYVGPLARGNSRFAWMTRKSEETITPWNFEEVVDKGASAQSFIERMTNFDKNLPNEKVL<br>PKHSLLYEYFTVYNELTKVKYVTEGMRKPAFLSGEQKKAIVDLLFKTNRKVTVKQLKEDYFKKIECFDSVEISG<br>VEDRFNASLGTYHDLLKIIKDKDFLDNEENEDILEDIVLTLTLFEDREMIEERLKTYAHLFDDKVMKQLKRRRYT<br>GWGRLSRKLINGIRDKQSGKTILDFLKSDGFANRNFMQLIHDDSLTFKEDIQKAQVSGQGDSLHEHIANLAGSPA |

IKKGILQTVKVVDELVKVMGRHKPENIVIEMARENQTTQKGQKNSRERMKRIEEGIKELGSQILKEHPVENTQLQ
NEKLYLYYLQNGRDMYVDQELDINRLSDYDVDAIVPQSFLKDDSIDNKVLTRSDKNRGKSDNVPSEEVVKKMK
NYWRQLLNAKLITQRKFDNLTKAERGGLSELDKAGFIKRQLVETRQITKHVAQILDSRMNTKYDENDKLIREVK
VITLKSKLVSDFRKDFQFYKVREINNYHHAHDAYLNAVVGTALIKKYPKLESEFVYGDYKVYDVRKMIAKSEQ
EIGKATAKYFFYSNIMNFFKTEITLANGEIRKRPLIETNGETGEIVWDKGRDFATVRKVLSMPQVNIVKKTEVQT
GGFSKESILPKRNSDKLIARKKDWDPKKYGGFDSPTVAYSVLVVAKVEKGKSKKLKSVKELLGITIMERSSFEK
NPIDFLEAKGYKEVKKDLIIKLPKYSLFELENGRKRMLASAGELQKGNELALPSKYVNFLYLASHYEKLKGSPED
NEQKQLFVEQHKHYLDEIIEQISEFSKRVILADANLDKVLSAYNKHRDKPIREQAENIIHLFTLTNLGAPAAFKYF
DTTIDRKRYTSTKEVLDATLIHQSITGLYETRIDLSQLGGDSGGSSGGSKRTADGSEFESPKKKRKVSGGSSGGST
LNIEDEYRLHETSKEPDVSLGSTWLSDFPQAWAETGGMGLAVRQAPLIIPLKATSTPVSIKQYPMSQEARLGIKP
HIQRLLDQGILVPCQSPWNTPLLPVKKPGTNDYRPVQDLREVNKRVEDIHPTVPNPYNLLSGLPPSHQWYTVLD
LKDAFFCLRLHPTSQPLFAFEWRDPEMGISGQLTWTRLPQGFKNSPTLFNEALHRDLADFRIQHPDLILLQYVDD
LLLAATSELDCQQGTRALLQTLGNLGYRASAKKAQICQKQVKYLGYLLKEGQRWLTEARKETVMGQPTPKTP
RQLREFLGKAGFCRLFIPGFAEMAAPLYPLTKPGTLFNWGPDQQKAYQEIKQALLTAPALGLPDLTKPFELFVDE
KQGYAKGVLTQKLGPWRRPVAYLSKKLDPVAAGWPPCLRMVAAIAVLTKDAGKLTMGQPLVILAPHAVEALV
KQPPDRWLSNARMTHYQALLLDTDRVQFGPVVALNPATLLPLPEEGLQHNCLDILAEAHGTRPDLTDQPLPDA
DHTWYTDGSSLLQEGQRKAGAAVTTETEVIWAKALPAGTSAQRAELIALTQALKMAEGKKLNVYTDSRYAFA
TAHIHGEIYRRRGWLTSEGKEIKNKDEILALLKALFLPKRLSIIHCPGHQKGHSAEARGNRMADQAARKAAITET
PDTSTLLIENSSPSGGSKRTADGSEFESPKKKRKVGSGPAAKRVKLD*

**Supplementary Note 2 |** Complete image of Western blot.

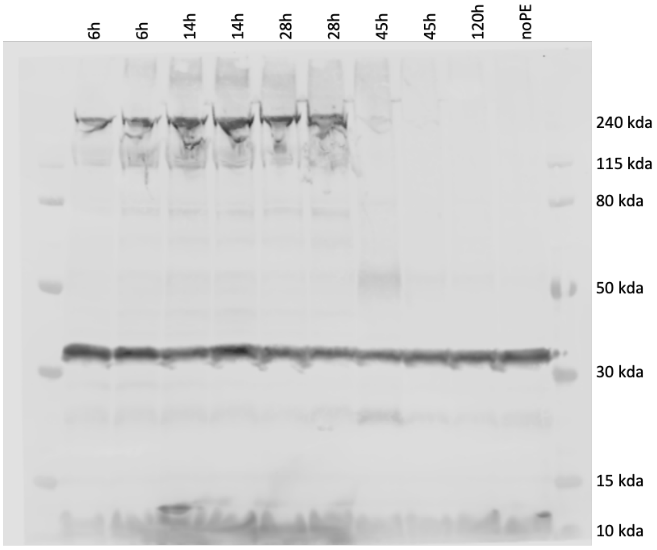


**Supplementary Table 1 |** pegRNA designs tested for correction of the *Pah*$^{enu}$ disease locus (5' > 3').

| | |
|---|---|
| peg-mPKU-2.1 | GCCTAATGTACTGTGTGCAG..scaffold..TTCCGGGTCTTCCACTGCACACAGTACA |
| peg-mPKU-2.1-tevo | GCCTAATGTACTGTGTGCAG..scaffold..TTCCGGGTCTTCCACTGCACACAGTACA-tevo |
| peg-mPKU-2.1-tmp | GCCTAATGTACTGTGTGCAG..scaffold..TTCCGGGTCTTCCACTGCACACAGTACA-tmp |
| peg-mPKU-0SM_tevo | GCCTAATGTACTGTGTGCAG..scaffold..TTCCGAGTCTTCCACTGCACACAGTACA-tevo |
| peg-mPKU-SM_tevo | GCCTAATGTACTGTGTGCAG..scaffold..TTCCGAGTCTTTCACTGCACACAGTACA-tevo |
| peg-mPKU_2SM_tevo | GCCTAATGTACTGTGTGCAG..scaffold..TTCCGAGTATTTCACTGCACACAGTACA-tevo |

**Supplementary Table 2 |** Oligonucleotides used for this study.

| | |
|---|---|
| HTS_PKU_fw | CTTTCCCTACACGACGCTCTTCCGATCTNNNNNNCCGTCCTGTTGCTGGCTTAC |
| HTS_PKU_rv | GGAGTTCAGACGTGTGCTCTTCCGATCTNNNNNNTGAGCATCCATTGTGGTTGG |
| peg-PKU_spacer_fw | ATGGTCTCGCACCGCCTAATGTACTGTGTGCAGGTTTCAGAGCTATGCTGGAAACAGC |
| peg-PKU-2.1_tevo_rv | ATGGTCTCGCGCGTGTACTGTGTGCAGTGGAAGACCCGGAAGCACCGACTCGGTGCCAC |
| peg-PKU-0SM_tevo_rv | ATGGTCTCGCGCGTGTACTGTGTGCAGTGGAAGACTCGGAAGCACCGACTCGGTGCCAC |
| peg-PKU-SM_tevo_rv | ATGGTCTCGCGCGTGTACTGTGTGCAGTGAAAGACTCGGAAGCACCGACTCGGTGCCAC |
| peg-PKU-2SM_tevo_rv | ATGGTCTCGCGCGTGTACTGTGTGCAGTGAAATACTCGGAAGCACCGACTCGGTGCCAC |
| PKU_CHANGE_1_fw | CTTTCCCTACACGACGCTCTTCCGATCTGGTTTCCCATTTCCCATCATCATT |
| PKU _CHANGE_1_rv | GGAGTTCAGACGTGTGCTCTTCCGATCTCTGGGAAGTGTGTACATGTATGG |
| PKU _CHANGE_2_fw | CTTTCCCTACACGACGCTCTTCCGATCTAGCATGTATGGGTTCCGAGG |
| PKU _CHANGE_2_rv | GGAGTTCAGACGTGTGCTCTTCCGATCTGAAACCAGTTTCAGCACGGTC |
| PKU _CHANGE_3_fw | CTTTCCCTACACGACGCTCTTCCGATCTATTAGGGAGGAGGGTAGAAGTGT |
| PKU _CHANGE_3_rv | GGAGTTCAGACGTGTGCTCTTCCGATCTGCAGTACATGAGCTTCCGC |
| PKU _CHANGE_4_fw | CTTTCCCTACACGACGCTCTTCCGATCTCAGGGCAAGCAGGTAGATGT |
| PKU _CHANGE_4_rv | GGAGTTCAGACGTGTGCTCTTCCGATCTACCTGTCGCCCTTCAGTTTG |
| PKU _CHANGE_5_fw | CTTTCCCTACACGACGCTCTTCCGATCTAGTATACCACTGTTTGTGCATTCC |
| PKU _CHANGE_5_rv | GGAGTTCAGACGTGTGCTCTTCCGATCTGCAAGTACGCTGCACACAAT |
| HTS_DNMT1_fw | CTTTCCCTACACGACGCTCTTCCGATCTNNNNNNGTCTTCCCCCACTCTCTTGC |
| HTS_DNMT1_rv | GGAGTTCAGACGTGTGCTCTTCCGATCTNNNNNNCCCCCAATATATGCCTCGGC |
| peg-DNMT1_spacer_fw | ATGGTCTCGCACCGCGGGCTGGAGCTGTTCGCGCGTTTCAGAGCTATGCTGGAAACAGC |
| peg-DNMT1_tevo_rv | ATGGTCTCGCGCGCTGGAGCTGTTCGCGCTGCCATCTTGCACCGACTCGGTGCCAC |
| DNMT1_CHANGE_1_fw | CTTTCCCTACACGACGCTCTTCCGATCTCTCAGCCGGACGCCCAATTA |
| DNMT1_CHANGE_1_rv | GGAGTTCAGACGTGTGCTCTTCCGATCTCCCGCAGCAGCCCTGT |
| DNMT1_CHANGE_2_fw | CTTTCCCTACACGACGCTCTTCCGATCTACAGGATGTGATATCGGAGGC |
| DNMT1_CHANGE_2_rv | GGAGTTCAGACGTGTGCTCTTCCGATCTCATGTGATCCACACACGCTT |
| DNMT1_CHANGE_3_fw | CTTTCCCTACACGACGCTCTTCCGATCTCGAAGGGGAAAACCCAGGAA |
| DNMT1_CHANGE_3_rv | GGAGTTCAGACGTGTGCTCTTCCGATCTGGTGGCACTGCTAGATCTCC |
| DNMT1_CHANGE_4_fw | CTTTCCCTACACGACGCTCTTCCGATCTAGCAACTTGAAAAGACTGTGCC |
| DNMT1_CHANGE_4_rv | GGAGTTCAGACGTGTGCTCTTCCGATCTAGGAGCTACGCAGAACCCTTC |
| DNMT1_CHANGE_5_fw | CTTTCCCTACACGACGCTCTTCCGATCTTCTCTTTTTGAATCTCCATAGCCCA |
| DNMT1_CHANGE_5_rv | GGAGTTCAGACGTGTGCTCTTCCGATCTTCACTTGTGAATGTAACACGGC |

**Supplementary Table 3** | Oligonucleotides used for RT-qPCR.

| | |
|---|---|
| SYBR_RT_Fw | GAAGGGCTGCAACACAACTG |
| SYBR_RT_Rev | AGCCCAGATTACCTCGGTCT |
| SYBR_Cas_Fw | ACCATCGACCGGAAGAGGTA |
| SYBR_Cas_Rev | TCGCTTGTTCCTGGTGTCTC |
| SYBR_mRPLP0_FW | TGAGATTCGGGATATGCTGTTGG |
| SYBR_mRPLP0_Rev | CGGGTCCTAGACCAGTGTTCT |

**Supplementary Table 4** | modified pegRNAs used in this study.

| | |
|---|---|
| **2’OMe-PKU** | mG*mC*mC*UAAUGUACUGUGUGCAGGUUUCAGAmGmCmUmAm<br>UmGmCmUmGmGmAmAmAmCmAmGmCmAmUmAmGmCAAGUU<br>GAAAUAAGGCUAGUCCGUUAUCAmAmCmUmUmGmAmAmAmAm<br>AmGmUmGmGmCmAmCmCmGmAmGmUmCmGmGmUmGmCUUC<br>CGAGUCUUUCACUGCACACAGUACAmU*mU*mA* |
| **2’OMe-PACE-PKU**<br>Tanja pegRNA 0922 | mG*mC*mC*UAAUGUACUGUGUGCAGGUUUCAGAGCUAUGCUGGA<br>AACAGCAUAGCAAGUUGAAAUAAGGCUAGUCCGUUAUCAACUUG<br>AAAAAGUGGCACCGAGUCGGUGCUUCCGAGUCUUUCACUGCACAC<br>AGUmA^mC^A |
| **2’OMe-scaffold-PACE-PKU**<br>PKU peg 0707 | mG*mC*mC*UAAUGUACUGUGUGCAGmGUUUUAGmAmGmCmUmA<br>mGmAmAmAmUmAmGmCmAmAGUUmAAmAAUAmAmGmGmCmUm<br>AGUmCmCGUUAmUmCAAmCmUmUmGmAmAmAmAmAmGmUmGGm<br>CmAmCmCmGmAmGmUmCmGmGmUmGmCUUCCGAGUCUUUCACU<br>GCACACAGUmA^mC^A |
| **2’OMe-scaffold-PACE-DNA-PKU**<br>**PKU peg DNA 0707** | mG*mC*mC*UAAUGUACUGUGUGCAGmGUUUUAGmAmGmCmUmA<br>mGmAmAmAmUmAmGmCmAmAGUUmAAmAAUAmAmGmGmCmUm<br>AGUmCmCGUUAmUmCAAmCmUmUmGmAmAmAmAmAmGmUmGGm<br>CmAmCmCmGmAmGmUmCmGmGmUmGmCdTdTdCdCdGdAdGdTdCdT<br>dTdTdCdAdCdTdGCACACAGUmA^mC^A |
| **2’OMe-scaffold-PACE -DNMT1**<br>DNMT1 peg 0707 | mC*mG*mG*GCUGGAGCUGUUCGCGCmGUUUUAGmAmGmCmUmA<br>mGmAmAmAmUmAmGmCmAmAGUUmAAmAAUAmAmGmGmCmUm<br>AGUmCmCGUUAmUmCAAmCmUmUmGmAmAmAmAmAmGmUmGGm<br>CmAmCmCmGmAmGmUmCmGmGmUmGmCAAGAUGGCAGCGCGAA<br>CAGCUCmC^mA^G |
| **2’OMe-scaffold-PACE-DNA-DNMT1**<br>DNMT1 peg DNA 0707 | mC*mG*mG*GCUGGAGCUGUUCGCGCmGUUUUAGmAmGmCmUmA<br>mGmAmAmAmUmAmGmCmAmAGUUmAAmAAUAmAmGmGmCmUm<br>AGUmCmCGUUAmUmCAAmCmUmUmGmAmAmAmAmAmGmUmGGm<br>CmAmCmCmGmAmGmUmCmGmGmUmGmCdAdAdGdAdTdGdGdCdAd<br>GdCdGCGAACAGCUCmC^mA^G |

| Modification Description | Notation when used in sequence |
|---|---|
| **2’O-methyl (2OMe)** | mA, mC, mG, mU |
| **2’-ribo 3’-phosphorothioate (S)** | A*, C*, G*, U* |
| **Deoxy (D)** | dA, dC, dG, dT |
| **2’-ribo 3’-phosphonoacetate (PACE)** | A^, C^, G^, U^ |
